# Hierarchized phosphotarget binding by the seven human 14-3-3 isoforms

**DOI:** 10.1101/2020.07.24.220376

**Authors:** Gergo Gogl, Kristina V. Tugaeva, Pascal Eberling, Camille Kostmann, Gilles Trave, Nikolai N. Sluchanko

**Affiliations:** Equipe Labellisee Ligue 2015, Department of Integrated Structural Biology, Institut de Genetique et de Biologie Moleculaire et Cellulaire (IGBMC), INSERM U1258/CNRS UMR 7104/Universite de Strasbourg, 1 rue Laurent Fries, BP 10142, F-67404 Illkirch, France; A.N. Bach Institute of Biochemistry, Federal Research Center of Biotechnology of the Russian Academy of Sciences, 119071, Moscow, Russia

**Author notes:** Corresponding authors: G.T.; N.N.S. Contributed equally: G.G. and K.V.T. Email addresses of the authors.

**Keywords:** 14-3-3 proteins, affinity trends, HPV E6 oncoprotein, fusicoccin, phosphopeptides, phosphorylation

## Abstract

The seven human 14-3-3 isoforms, highly similar yet encoded by distinct genes, are among the top 1% highest-expressed human proteins. 14-3-3 proteins recognize phosphorylated motifs within numerous human or viral proteins. We analyzed by crystallography, fluorescence polarization, mutagenesis and fusicoccin-mediated modulation the structural basis and druggability of 14-3-3 binding to four E6 oncoproteins of tumorigenic HPV. The seven isoforms bound variant and mutated phospho-motifs of E6 and unrelated protein RSK1 with different affinities, albeit following an ordered ranking profile with conserved relative K_D_ ratios. Remarkably, 14-3-3 isoforms obey the same hierarchy when binding to most of their established targets, nicely supported by a recent proteome-wide human complexome map. This knowledge allows predicting the proportions of 14-3-3 isoforms engaged with phosphoproteins in various tissues. Notwithstanding their individual functions, cellular concentrations of 14-3-3 may be collectively adjusted to buffer the strongest phosphorylation outbursts, explaining their expression variations in different tissues and tumors.

## INTRODUCTION

14-3-3 proteins recognize protein partners phosphorylated at serine or threonine in certain sequence motifs in all eukaryotic organisms. The seven human 14-3-3 “isoforms”, individually named β, γ, ε, ζ, η, σ, and τ (beta, gamma, epsilon, zeta, eta, sigma and tau) ^1^, are distinct gene encoded paralogs which are highly similar in sequence and in their phosphopeptide-recognition mode, yet display different expression patterns across tissues ^2, 3^. 14-3-3 proteins are highly abundant in most human tissues, where several 14-3-3 isoforms are systematically found among the top 1% of the ∼20,000 human gene-encoded proteins ^3^. For instance, according to the Protein Abundance Database, PAXdb, ^3^ the cumulated seven 14-3-3 isoforms are within the five most abundant protein species in platelets.

14-3-3 proteins function as dimers able to bind phosphopeptides ^4, 5^. Phosphorylated 14-3-3-binding sequences usually correspond to internal motifs I RSX(pS/pT)X(P/G) and II RXY/FX(pS/pT)X(P/G) ^5^ and to the C-terminal motif III (pS/pT)X_0-2_-COOH ^6, 7^, where pS/pT denotes phosphorylated serine or threonine and X denotes any amino acid. The regulation by 14-3-3 binding typically protects 14-3-3 targets from dephosphorylation, thereby affecting their activities, their interactions with other proteins, their turnover and intracellular localization ^8^. 14-3-3 proteins are indispensable in a diversity of processes such as apoptosis, cell cycle, or signal transduction ^1, 9^. They are involved in neurodegenerative disorders, viral infection and cancer, often representing promising drug targets ^10^.

14-3-3 also directly interact with several viral proteins ^11^, such as the E6 oncoprotein of high-risk mucosal human papillomaviruses (hrm-HPV) ^12, 13, 14^ responsible for genital cancers (cervix, anus) and a growing number of head-and-neck cancers ^15, 16^. E6 is one of the two main early-expressed HPV oncoproteins. In HPV-transformed cells, E6 interacts with numerous host proteins ^17^ to counteract apoptosis, alter differentiation pathways, polarity and adhesion properties and thereby sustain cell proliferation ^18, 19^. Inhibition of E6 in HPV-positive cell lines results in the cell growth arrest and induces apoptosis or rapid senescence ^20, 21, 22, 23^. All hrm-HPV E6 proteins harbor a phosphorylatable dual-specificity C-terminal motif ^24^ (Fig. 1A). In its unphosphorylated state, it is a PDZ-domain binding motif (PBM) that mediates E6 binding to a range of cognate host proteins regulating cell polarity, adhesion, differentiation or survival ^14^. When the motif is phosphorylated, E6 proteins, in particular those of hrm-HPV 16, 18 and 31, acquire the capacity to bind 14-3-3 ^12, 13, 25^.

**Fig. 1.**
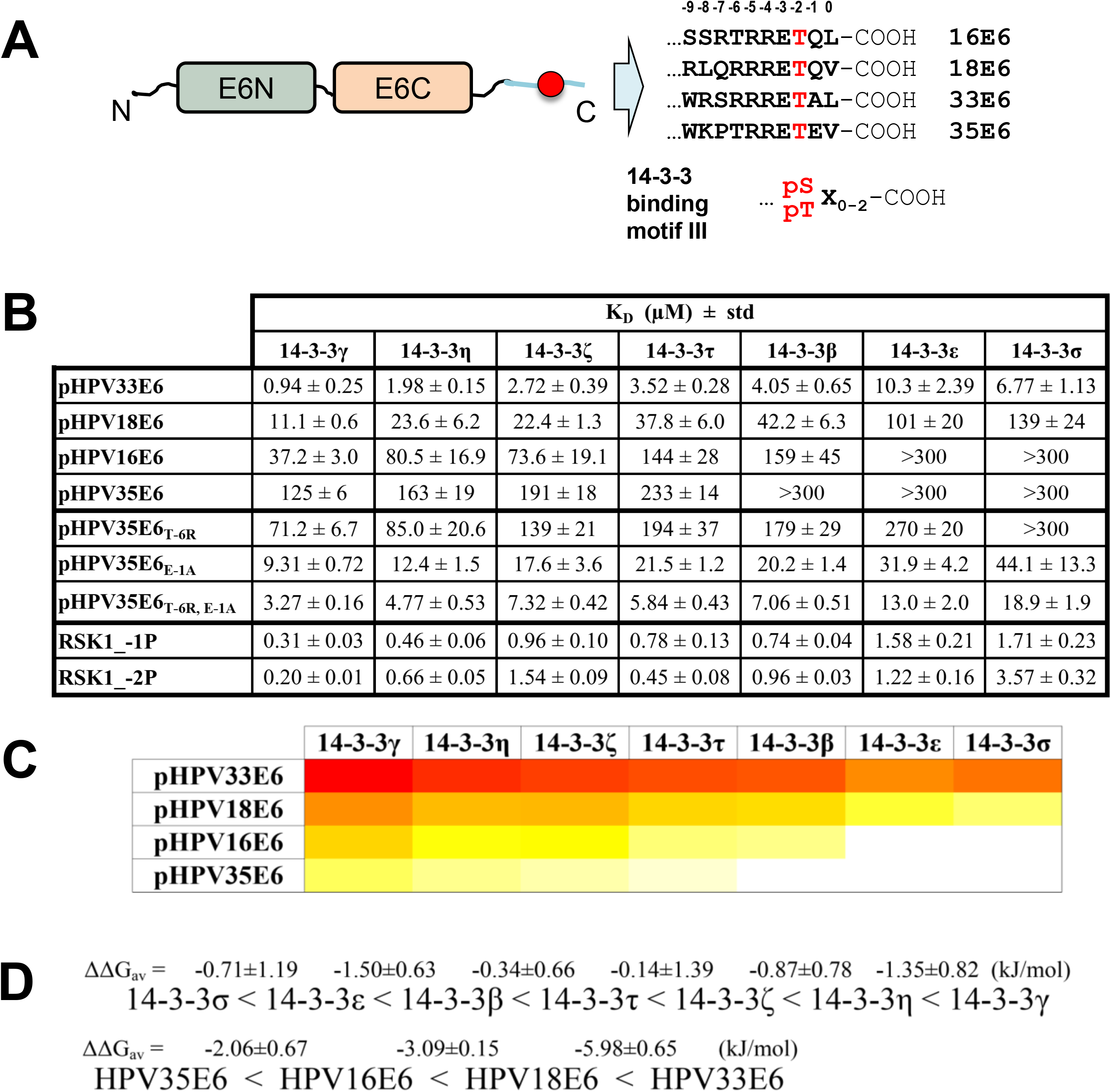
Selected E6 PBMs reveal parallel binding profiles to human 14-3-3 isoforms. **A**. Exemplary phosphorylatable C-terminal E6 PBMs from high-risk mucosal HPV types overlap with the 14-3-3-binding motif III ^6^. The positions are numbered above, according to conventional PBM numbering. **B**. Affinities of four selected HPV-E6 phospho-PBMs, p35E6 mutants and RSK1 phosphopeptides towards the seven human 14-3-3 isoforms as determined by fluorescence polarization using FITC-labeled HSPB6 phosphopeptide as a tracer. Apparent K_D_ values determined from competitive FP experiments are presented. **C**. The heatmap representation of the data on panel A showing the affinity trends in the interaction profiles between 14-3-3 isoforms and four HPV-E6 phospho-PBMs from weakest (white) to strongest (red). **D**. Averaged ΔΔG values between 14-3-3 isoforms or E6 phospho-PBM pairs, calculated based on their observed order of binding affinities (from weakest to strongest). Individual K_D_ values from Supplementary Fig. 2 were first converted into ΔG values (at T=295 K; excluding cases when K_D_ > 300 μM) and average ΔΔG values (ΔΔG_av_) were calculated between the indicated motifs/isoforms.

Here we study the structural basis and druggability of 14-3-3 binding to E6 oncoproteins of four tumorigenic HPV types by a combination of crystallography, binding assays, and mutagenesis. We show that the seven isoforms bound phospho-PBMs of E6 proteins and of the unrelated human RSK kinase with different affinities, albeit obeying a hierarchized profile with conserved relative K_D_ ratios. This hierarchy turns out to be a general feature of the interaction of 14-3-3 isoforms with most of their targets, supported by both literature and a recently released proteome-wide human complexome map ^26^. Using this knowledge, we built a predictor that estimates the proportions of 14-3-3 isoforms engaged with phosphoproteins in various human tissues, cell lines or tumors.

## RESULTS

### Selected E6 PBMs reveal parallel binding profiles to human 14-3-3 isoforms

Among all 225 HPV E6 proteins curated in the PaVE database (https://pave.niaid.nih.gov/, last accessed on 28 September 2020), 31 E6 proteins from mucosal α-genera HPV possess a C-terminal PBM with the class 1 consensus (X(S/T)X(L/V/I/C)-COOH, where X is any amino acid residue ^27, 28^). E6 PBMs are phosphorylatable by protein kinases at their conserved antepenultimate S/T residue ^12, 13, 29^. This phosphosite is preceded by arginine residues in most of the HPV-E6 PBM sequences with recognizable basophilic kinase substrate consensus motifs, R(X/R)X(S/T) and RXRXX(S/T) ^30, 31^. The E6 PBMs can be classified in three subgroups: subgroups 1 and 2 prone to phosphorylation by the basophilic kinases and “orphan” subgroup 3 with a less predictable phosphorylation propensity (Supplementary Fig. 1). In line with earlier observations ^12, 25, 32^, the phospho-PBM sequences from subgroups 1 and 2 ideally match the C-terminal 14-3-3-binding motif III ^6^ (Fig. 1A).

Four phospho-PBMs from E6 proteins of HPV types 16, 18, 33 and 35 belonging to subgroups 1 and 2 (as defined in Supplementary Fig. 1) were analyzed for their interaction with all seven full-length human 14-3-3 isoforms. For comparison, we also measured two non-viral phospho-PBMs originating from protein kinase RSK1 ^25^. We used a competitive fluorescence polarization assay that measures the displacement of a fluorescent tracer phosphopeptide (here, HSPB6) bound beforehand to 14-3-3, by an increasing amount of the peptide of interest. All binding curves are shown in Supplementary Fig. 2A.

All phospho-PBMs (p16E6, p18E6, p33E6, p35E6, RSK1_-1P, and RSK1_-2P) detectably bound to 14-3-3 proteins, in sharp contrast to their unphosphorylated counterparts. The interactions between E6 phospho-PBMs and 14-3-3 proteins spanned very wide affinity ranges, from just below 1 μM (p33E6–14-3-3γ) to above 300 μM (Fig. 1B and Supplementary Fig. 2A). Such large binding affinity differences are noteworthy since the four E6 PBM sequences are very similar (Fig. 1A), and all 14-3-3 isoforms share highly conserved phosphopeptide-binding grooves.

Remarkably, the six phospho-PBMs obeyed a consistent hierarchized profile in their relative binding preferences towards the seven 14-3-3 isoforms, albeit with an overall shift in affinity from one peptide to another. For each phosphopeptide, the seven 14-3-3 isoforms systematically clustered as four groups of decreasing affinity, in a conserved order from the strongest to the weakest phospho-PBM binder: gamma, eta, zeta/tau/beta and epsilon/sigma (γ, η, ζ /τ/β, and ε/σ) (Fig. 1C). These conserved relative affinity shifts can be quantified by calculating, for two distinct 14-3-3 isoforms, their differences of free energy of binding (ΔΔG) towards each individual phosphopeptide, then calculating the average difference (ΔΔG_av_) with its standard deviation (Fig. 1D). Between the strongest and the weakest binders (isoforms γ and σ, respectively) the average phosphopeptide-binding energy difference is ΔΔG_av_ = −5.1 ± 1.3 kJ/mol, roughly corresponding to a 11-fold K_D_ ratio.

The seven 14-3-3 isoforms also showed consistent profiles in their relative binding preferences towards the four E6 phospho-PBMs. For each 14-3-3 isoform, the four phospho-PBMs systematically rank the same way from the strongest to the weakest binder: p33E6, p18E6, p16E6 and p35E6 (Fig. 1D). The average 14-3-3 binding free energy difference between p33E6 and p35E6 was ΔΔG_av_ = −10.9 ± 0.7 kJ/mol, roughly corresponding to a 100-fold K_D_ ratio.

### Atomic structure reveals the 14-3-3ζ –18E6 PBM interface

To get structural insight into the 14-3-3ζ interaction with 18E6 PBM, we determined a crystal structure of the 14-3-3ζ –18E6 phospho-PBM complex at a 1.9 Å resolution using a previously reported chimeric fusion strategy ^33, 34^ (Table 1, Fig. 2, Supplementary Figs 3 and 4). The phosphopeptide establishes multiple polar interactions with the basic pocket in the amphipathic groove of 14-3-3 (Supplementary Fig. 5), largely reminiscent of previously solved structures of 14-3-3–phosphopeptide complexes ^5, 33^. The conformation of 18E6 phosphopeptide bound to 14-3-3ζ within the chimera is practically identical (RMSD = 0.17 Å upon superimposition of Cα atoms of the peptides) to the 14-3-3σ-bound conformation of a synthetic 16E6 phosphopeptide reported very recently at a lower resolution (Fig. 2B) ^25^. The observed conservation of most interface contacts within the two complexes suggest that these crystal structures can serve as templates to build accurate homology models of 14-3-3 complexes for other E6 phospho-PBMs or, more generally, other C-terminal motif III peptides phosphorylated on the antepenultimate position.

**Fig. 2.**
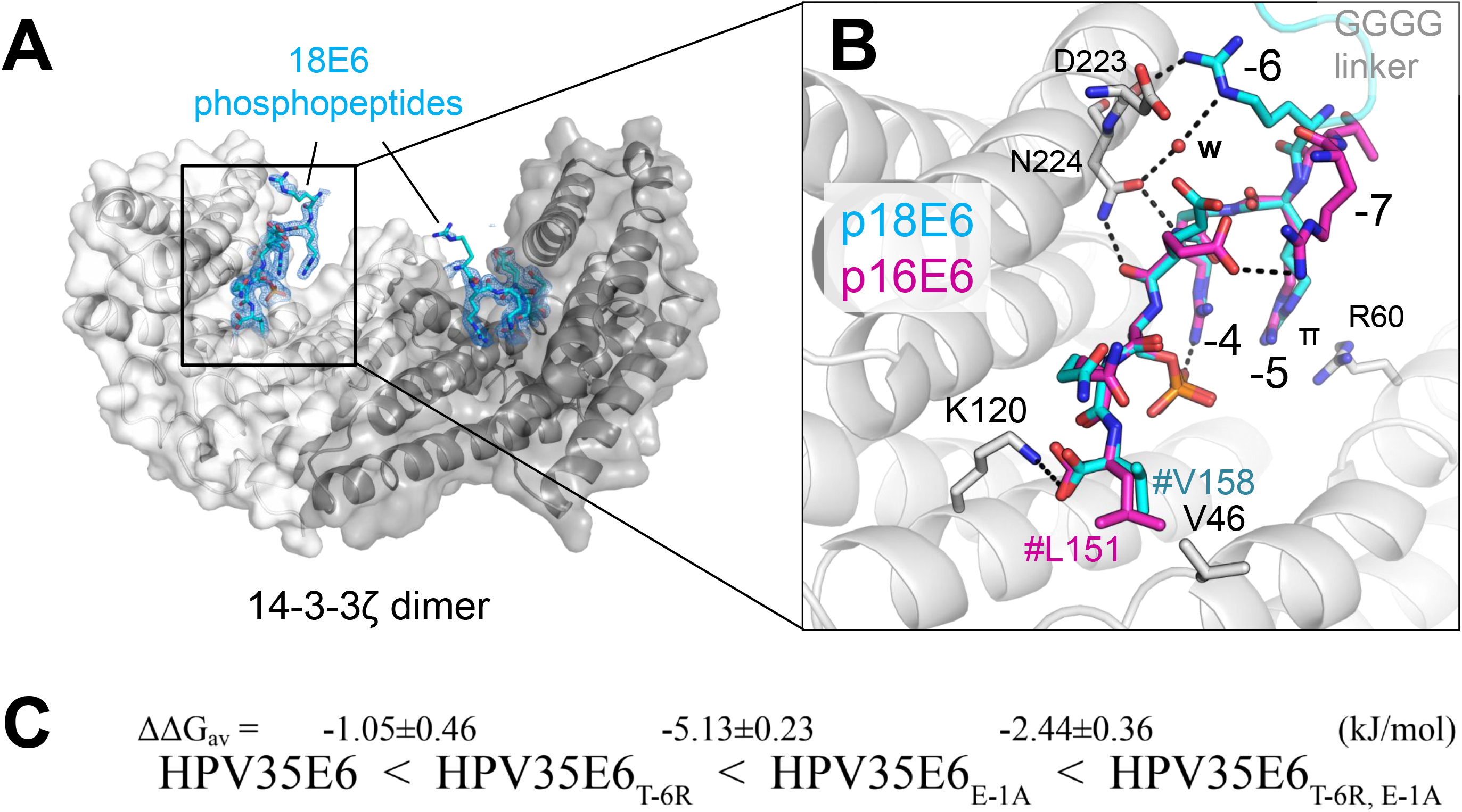
Molecular interface between 14-3-3ζ and phosphorylated 18E6 PBM at a 1.9 Å resolution. **A**. An overall view on the 14-3-3ζ dimer (subunits are in tints of grey) with two bound 18E6 phosphopeptides (cyan sticks). **B**. An overlay of the two 14-3-3 bound phosphopeptides from 16E6 (6TWZ.pdb) and 18E6 (this work) showing the similarity of the conformation. # denotes the C-terminus (-COOH). w – the water molecule, π – π-stacking interaction. Key positions are numbered according to the PBM convention. **C**. Averaged ΔΔG values between 14-3-3 isoforms or 35E6 phospho-PBM pairs, calculated based on their observed order of binding affinities (from weakest to strongest). Individual K_D_ values from Supplementary Fig. 2 were first converted into ΔG values (at T=295 K; excluding cases when K_D_ > 300 μM) and average ΔΔG values (ΔΔG_av_) were calculated between the indicated motifs/isoforms.

**Table 1.** Crystallographic statistics.

|  | **14-3-3ζ-18E6 chimera** | **14-3-3ζ-18E6 chimera + FSC** |
| --- | --- | --- |
| ***Data collection*** |  |  |
| Wavelength | 1.00 | 1.00 |
| Resolution range | **39.26 - 1.9** (1.95 - 1.9) | **38.05 - 1.85** (1.9 - 1.85) |
| Space group | P 21 21 21 | P 21 21 21 |
| Unit cell (a, b, c, α, β, γ) | 72.35, 78.53, 90.3, 90, 90, 90 | 73.23, 76.1, 88.95, 90, 90, 90 |
| Total reflections | 547783 (37726) | 557315 (41375) |
| Unique reflections | 41285 (2986) | 41927 (3038) |
| Multiplicity | 13.3 (12.6) | 13.3 (13.6) |
| Completeness (%) | 100 (100) | 97.1 (96.5) |
| Mean I/sigma(I) | 13.19 (1.39) | 12.36 (1.40) |
| R-meas | 16.6 (205) | 16.3 (216) |
| CC1/2 | 99.9 (54.7) | 99.8 (58.4) |
| ***Refinement*** |  |  |
| R-work | 0.1764 | 0.1908 |
| R-free | 0.2071 | 0.2192 |
| Number of non-hydrogen atoms | 4468 | 4589 |
| macromolecules | 4034 | 4004 |
| ligands | 24 | 102 |
| solvent | 410 | 483 |
| Protein residues | 481 | 482 |
| RMS(bonds) | 0.006 | 0.006 |
| RMS(angles) | 0.83 | 0.71 |
| Ramachandran favored (%) | 99.36 | 98.51 |
| Ramachandran allowed (%) | 0.64 | 1.49 |
| Ramachandran outliers (%) | 0 | 0 |
| Rotamer outliers (%) | 1.65 | 2.13 |
| Clashscore | 6.68 | 3.17 |
| Average B-factor | 33.39 | 29.6 |
| macromolecules | 32.42 | 28.49 |
| ligands | 62.93 | 32.67 |
| solvent | 41.2 | 38.11 |
| Number of TLS groups | 15 | 11 |
| PDB ID | **6ZFD** | **6ZFG** |

Nonetheless, a few noteworthy differences appear in a subset of the crystallographic conformers of 14-3-3/16E6 and 14-3-3/18E6 complexes. On the one hand, in 1 of the 4 conformers observed in the asymmetric unit of the 14-3-3σ/16E6 crystal, the side chains of Arg −7 (Gln in 18E6) and Glu −3 form an additional *in-cis* salt bridge (Fig. 2B). On the other hand, Arg −6 of 18E6 (Thr in 16E6) mediates a bipartite interaction with 14-3-3 in most of the observed conformers. It simultaneously interacts with the carbonyl of Asp223 and participates in a water-mediated interaction with Asn224 (Fig. 2B and Supplementary Fig. 5).

### Rescue of the weakest E6–14-3-3 interaction by rational design

Next, we investigated possible causes of the remarkable 14-3-3 binding affinity differences observed between the four E6 phospho-PBMs.

In principle, the affinity of a series of variant peptides for a given protein may be modulated by two types of atomic contacts: intermolecular and intramolecular contacts within the formed complexes, and intramolecular contacts in the free unbound peptides.

As concerns contacts within the 14-3-3/E6 complexes, the crystal structures have shown that Arg −6 can mediate more interactions than Thr −6 with the generic 14-3-3 interface (Fig. 2B). Interestingly, position −6 is an Arg in the two strongest 14-3-3-binders (18E6 and 33E6) versus a Thr in the weakest ones (35E6 and 16E6).

As concerns possible contacts within the unbound peptides, we noticed that all E6 phospho-PBMs have a delicate charge distribution, with an acidic C-terminal segment (that includes the C-terminus and the natural acidic or phosphorylated residues) and a basic N-terminal segment (that is also involved in recognition by kinases). These local charged segments may form transient *in-cis* interactions within the unbound phosphopeptide, so-called “charge clamps” ^35^. We speculated that Glu −1 in p35E6, the weakest 14-3-3 binder, might participate in such a charge clamp, thereby disfavoring its binding to 14-3-3.

To address these potential mechanisms, we synthesized three variants of the weakest 14-3-3 binder, p35E6. The first variant contained a T-6R substitution, which in principle could allow a more stable bound conformation, but may also stabilize charge clamps in the free form of the motif. The second variant contained an E-1A substitution, which in principle could destabilize *in-cis* charge-clamps. A third variant contained both substitutions. All substitutions turned out to reinforce the binding affinities of 35E6 without altering the apparent preferences of the different 14-3-3 isoforms (Fig. 1B and Fig. 2C). Taken individually, T-6R moderately increased binding (ΔΔG_av_ = −1.1 ± 0.5 kJ/mol, 1.5-fold K_D_ ratio), while E-1A strongly reinforced it (ΔΔG_av_ = −5.1 ± 0.2 kJ/mol, 11-fold K_D_ ratio). When combined, the two substitutions synergistically increased binding (ΔΔG_av_ = −8.7 ± 0.4 kJ/mol, 35-fold K_D_ ratio), thereby turning p35E6 into a strong 14-3-3 binder, just below p33E6. These results indicate that the two above-stated mechanisms act in combination to generate the wide 14-3-3-binding affinity range displayed by distinct E6 phospho-PBMs despite of their high sequence conservation.

### The 14-3-3/E6 PBM interaction is druggable by fusicoccin

Fusicoccin (FSC) is a commonly used stabilizer of 14-3-3 complexes, when its binding in the distinct pocket in the 14-3-3/phosphopeptide interface is allowed by phosphopeptide side chains of the amino acids in downstream positions relative to the phospho-residue ^36, 37, 38, 39^. This is especially the case with motif III phosphopeptide complexes of 14-3-3 having only one residue after the phosphosite ^37, 39, 40^. However, the effect of FSC on interaction of longer motif III phosphopeptides with 14-3-3 is less characterized (Supplementary Table 1).

We performed FP experiments to measure equilibrium binding affinity constants of complexes between the four HPV-E6 phosphopeptides and 14-3-3 isoforms ζ and γ, in the presence of 100 µM FSC (Supplementary Fig. 2B and Fig. 3A). The addition of FSC consistently decreased by 1.5 to 2 fold the affinities of all eight interactions (ΔΔG_av_ = −1.3 ± 0.5 and −1.8 ± 0.4 kJ/mol for ζ and γ, respectively) without altering the apparent preferences of the different peptides (Fig. 3A and B).

**Fig. 3.**
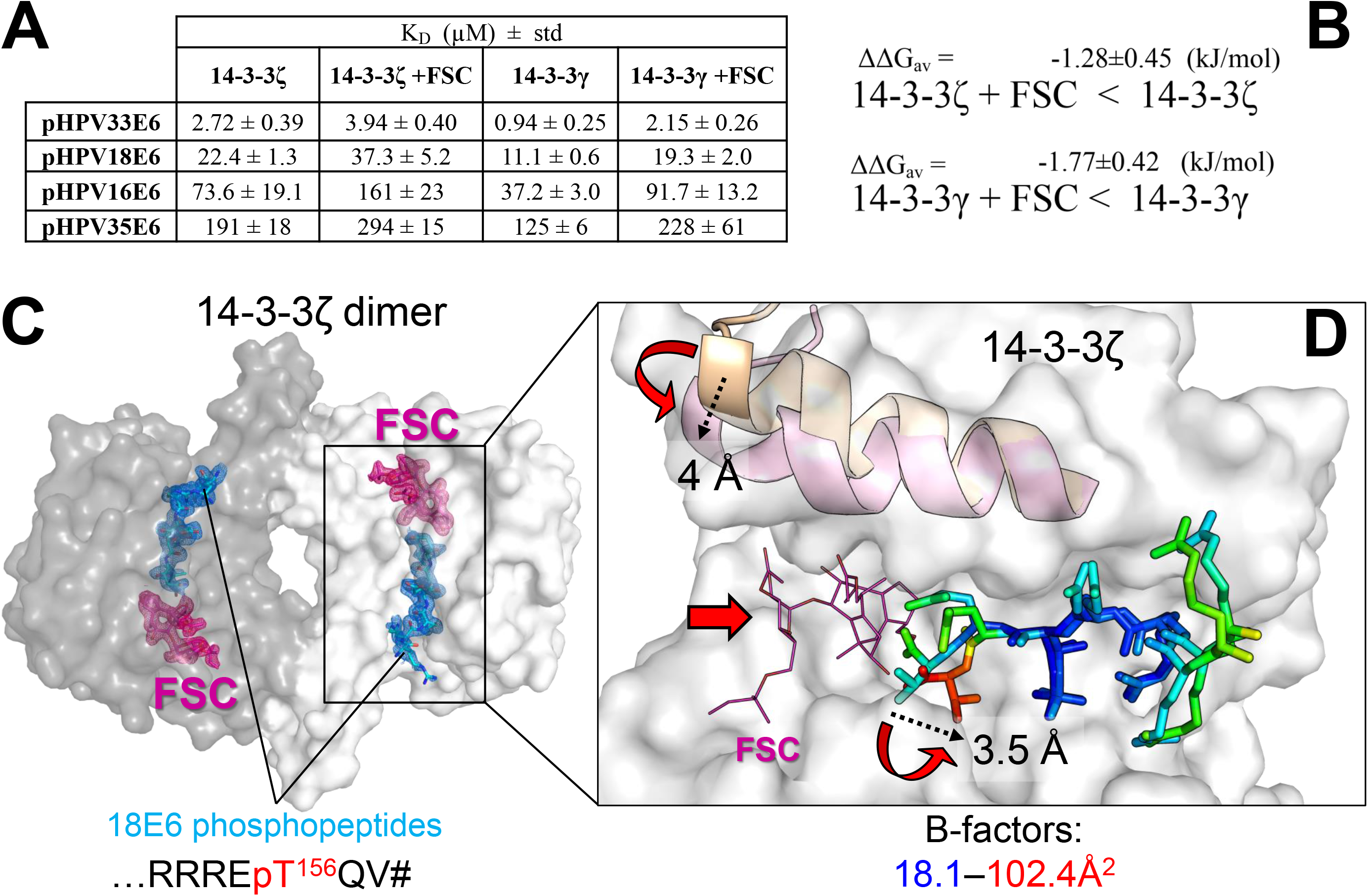
The 14-3-3ζ /18E6 PBM interaction is druggable by FSC. **A**. Affinities of four selected HPV-E6 phospho-PBMs towards human 14-3-3ζ and 14-3-3γ in the absence and in the presence of FSC as determined by fluorescence polarization using FITC-labeled HSPB6 phosphopeptide as a tracer. Apparent K_D_ values determined from competitive FP experiments are presented. The binding curves are shown in Supplementary Fig. 2. **B**. Averaged ΔΔG values between 14-3-3–E6 phospho-PBM pairs in the absence or in the presence of FSC, calculated based on their observed order of binding affinities (from weakest to strongest). Individual K_D_ values from Supplementary Fig. 2 were first converted into ΔG values (at T=295 K; excluding cases when K_D_ > 300 μM) and average ΔΔG values (ΔΔG_av_) were calculated between the indicated motifs/isoforms. **C**. An overall view on the ternary complex between 14-3-3ζ (subunits are shown by surface using two tints of grey), 18E6 phosphopeptide (cyan sticks) and FSC (pink sticks). FSC was soaked into the 14-3-3ζ –18E6 chimera crystals. 2F_o_-F_c_ electron density maps contoured at 1σ show are shown for the phosphopeptide and FSC only. **D**. The effect of FSC binding. Conformational changes upon FSC binding are shown by red arrows, a significant rise of the local B-factors of the phosphopeptide is shown using a gradient from blue to red as indicated. The amplitudes of the conformational changes in 14-3-3 and 18E6 peptide are indicated in Å by dashed arrows.

Next, we used a soaking approach to crystallize the ternary 14-3-3ζ /18E6 PBM/FSC complex and solved its structure at 1.85 Å resolution (Fig. 3C, Supplementary Fig. 4, 6 and Table 1). FSC binding in the well-defined cavity did not disrupt the overall assembly (Supplementary Fig. 4 and Fig. 3C), but it induced a hallmark ∼4 Å closure of the last α-helix of 14-3-3ζ (Fig. 3D) as observed for other 14-3-3 complexes containing FSC ^41^. Also, FSC binding reoriented the C-terminal carboxyl group and caused local destabilization of the very C-terminal residues of the phosphopeptide, increasing their temperature factors and dispersing the local electron density (Fig. 3D and Supplementary Fig. 6). As a result of FSC binding, the water network around the phospho-PBM C-terminus significantly changed (Supplementary Fig. 7).

Nevertheless, the simultaneous binding of FSC and E6 PBM in the amphipathic groove of 14-3-3 indicates that such ternary complex can be used as a starting point to design both stabilizers and inhibitors of 14-3-3/E6 interactions.

### Hierarchized peptide-affinity profiles are a general feature of human 14-3-3 isoforms

Former studies have measured the binding of the seven human 14-3-3 isoforms to unrelated phospho-motifs derived from Cystic Fibrosis Transmembrane Conductance Regulator (CFTR), Leucine-Rich Repeat Kinase 2 (LRRK2), Potassium channel subfamily K members (TASK1/3), C-Raf, the p65 subunit of the NF-κB transcription factor, and from Ubiquitin carboxyl-terminal hydrolase 8 (USP8), representing a wide variety of different 14-3-3-binding motifs, including C-terminal, internal, monovalent or divalent motifs ^38, 42, 43, 44, 45, 46^. These phosphorylated motifs from different origins have a strikingly wide affinity range, spanning from low nanomolar to low millimolar detectable dissociation constants (Fig. 4A-C). For instance, for 14-3-3γ, the K_D_ ratio between the strongest and the weakest binding phosphopeptide is almost 625-fold in the present work, and 39,000-fold when taking into account affinities from the literature (Fig. 4B).

**Fig. 4.**
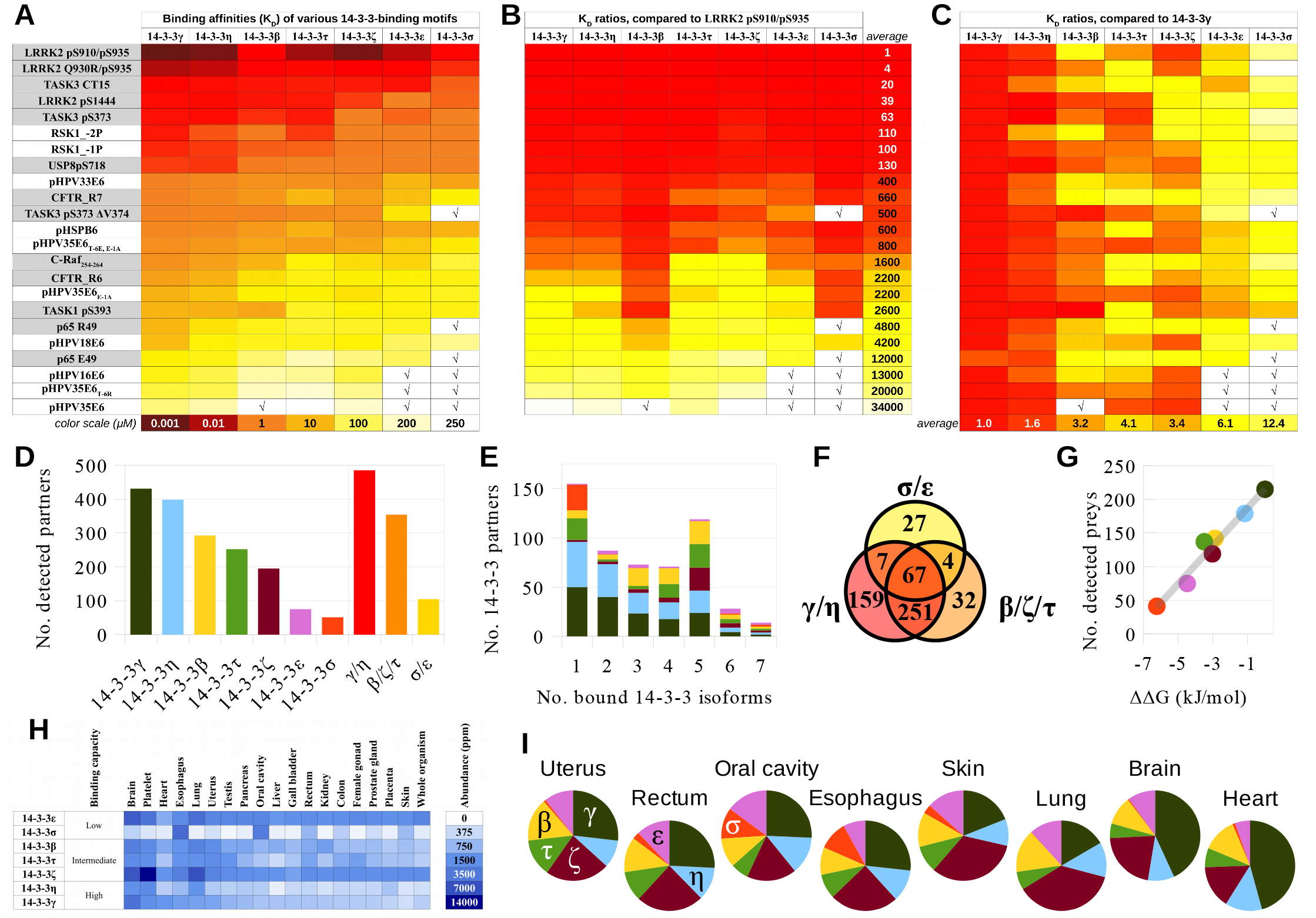
Hierarchized target binding by the seven human 14-3-3 isoforms is a general trend. **A**. Affinity maps of 14-3-3 interactions based on experimentally determined dissociation constants against the 14-3-3ome, as obtained in the current work and in ^38, 42, 43, 44, 45, 46^. Binding motifs that are analyzed in other studies are highlighted with a grey background ^38, 42, 43, 44, 45, 46^. The color scale is either based on affinity values or in K_D_ ratios. √ denotes affinities weaker than the limit of quantitation of the experimental assays. **B**. Same map as in (A), normalized to the strongest 14-3-3-binding motif. An average 34,000-fold K_D_ ratio is observed between the strongest and weakest 14-3-3-binding peptide. **C**. Same map as in (A), normalized to the strongest phosphopeptide-binder 14-3-3γ. Note that all peptides are following very similar affinity trends between the different 14-3-3 isoforms, with an average 12-fold K_D_ ratio between the strongest-binding and weakest-binding 14-3-3 isoform. **D**. Number of unique partners detected according to the Bioplex database (https://bioplex.hms.harvard.edu and ^26^) for each 14-3-3 isoform, taken individually (left) or grouped in three subsets (right) following their relative affinity trends (strong, intermediate, and weak binders). **E**. From left to right: number of 14-3-3 partners in the Bioplex database, that bound to 1, 2, 3, 4, 5, 6 or all 7 isoforms, respectively. Within each bar, the proportion of partners that bound to each individual isoform is indicated (same isoform color code as in (D)). **F**. Venn diagram showing the repartition of the 14-3-3 partners from the Bioplex database among the strong, medium and weak phosphopeptide-binding subsets, defined as in (D). **G**. Correlation between the number of “prey” binders of 14-3-3 isoforms used as “baits”, according to the Bioplex database, and the average free binding energy difference ΔΔG between the strongest phosphopeptide-binder 14-3-3γ, and all individual isoforms (same color code as in (D)). ΔΔG values were calculated from the average K_D_ ratios from panel C. **H**. Abundance of the seven 14-3-3 isoforms across different human tissues and in the whole human organism, according to the PAXdb database (https://pax-db.org and ^3^). **I**. Predicted proportions of 14-3-3-bound phosphoproteins that would be engaged with each individual isoform in different tissues, assuming that the majority of 14-3-3 molecules are available for interaction (same color code as in (D)).

Conversely, the hierarchized relative binding profile of the seven human 14-3-3 isoforms observed herein for E6 and RSK1 phosphopeptides is remarkably confirmed in most published data that have also measured affinities for all these seven isoforms ^38, 42, 43, 44, 45, 46^, with 14-3-3γ and 14-3-3η consistently being the strongest binders and 14-3-3σ and 14-3-3ε being the weakest binders, independently of the nature of the target motif (Fig. 4C). Furthermore, the average maximal K_D_ ratio between the strongest-binding and the weakest-binding 14-3-3 in the literature is around 12-fold, like in our present work (∼11-fold).

Moving further, we wondered whether the observed trends would be conserved at a full proteome-wide scale. The Gygi laboratory (Harvard University) applied a massive parallel affinity-purification coupled to mass-spectrometry (AP-MS) technique to decipher the complexomes of more than 10,000 recombinantly expressed bait proteins in two orthogonal cell lines ^26^. We retrieved from the “Bioplex3” database (a compendium of all data from the Gygi laboratory) the numbers of detected interaction partners for each 14-3-3 isoform (Fig. 4D-G). In total, 547 unique proteins were detected as an interaction partner of at least a single 14-3-3 isoform. Out of those, 14-3-3γ and 14-3-3η had the highest number of interaction partners, followed by a second group including 14-3-3β, 14-3-3ζ, and 14-3-3τ, and a third group comprising 14-3-3σ and 14-3-3ε (Fig. 4D). Most of these interaction partners were found to bind more than a single 14-3-3 isoform (Fig. 4F and G). While the strongest-binding isoforms (γ and η) do not share ∼30% of their interactome with the other isoforms, they interact with more than 85% of the binders of the mild-binding isoforms (β, ζ, and τ) and more than 90% of the binders of the weak-binding isoform 14-3-3ε. Indeed, out of the 75 detected binders of 14-3-3ε, only 1 (below 2% of the total) is unique to 14-3-3ε. By contrast, the other weak-binding isoform, 14-3-3σ, has a distinct behavior. Out of its 51 detected binders, 26 interactions are unique to 14-3-3σ (above 50% of all its binders).

In the AP-MS experiments, interaction partners (and 14-3-3 proteins in particular) can be either “baits” or “preys”. Baits are recombinantly expressed in the cells using the same promoter, which should ensure a relatively even expression for all 14-3-3 isoforms. By contrast, the preys are proteins naturally expressed by the cells, so that the distinct 14-3-3 preys should be present in very different amounts, depending on their intrinsic levels of expression in the host cells. We observed a remarkable linear correlation (R^2^ = 0.96) between the numbers of detected interaction partners (as preys) captured by different 14-3-3 isoforms (as baits) and their relative affinity (ΔΔG_av_) for the best phosphopeptide-binder, 14-3-3γ isoform (Fig. 4G). The correlation decreased when using 14-3-3 prey-binding baits (R^2^ = 0.84) or the using the mixture from 14-3-3 bait and 14-3-3 prey pools (R^2^ = 0.91) (Supplementary Fig. 8A). In support of these interrelations, the number of 14-3-3 prey-binding baits also indicated correlation with the affinity trend of 14-3-3 isoforms when using data from a recent independent study (https://sec-explorer.shinyapps.io/Kinome_interactions/ and ^47^) that used AP-MS to uncover the interactions of more than 300 protein kinases (R^2^ = 0.64) (Supplementary Fig. 8A).

Remarkably, the level of overall sequence divergence of 14-3-3 isoforms, using 14-3-3γ as a reference (γ < η < β ≈ ζ < τ < σ < ε; i.e., ε is the most divergent from γ; Supplementary Fig. 9A), also correlates very well with their hierarchized affinity differences (Fig. 1 and 4). However, the latter cannot be explained merely by features of the phosphopeptide-binding regions of 14-3-3 isoforms, which in fact are identical in all seven human 14-3-3 proteins (Supplementary Fig. 9A and B). Indeed, even the extreme isoforms on the peptide-affinity scale, 14-3-3γ and 14-3-3σ, have only minor sequence variations and only at the periphery of the peptide-binding grooves (Supplementary Fig. 9C), which are unlikely to dictate the phosphopeptide binding differences. Interestingly, the sequence divergence trend relative to 14-3-3γ (Supplementary Fig. 9A) remains conserved when considering diverse sub-regions of the sequence (Supplementary Fig. 10). This indicates that the general target affinity differences arise from fine conformational effects spanning the entire structure, rather than a defined sub-region.

### Prediction of cellular 14-3-3/phosphotargets complexomes

14-3-3 proteins are highly expressed. Therefore, their abundances in all human tissues have been reliably quantified. According to the integrated whole human body dataset of the Protein Abundance Database, PAXdb (https://pax-db.org and ^3^), 14-3-3ε is the 48th most abundant human protein (2479 ppm) and 14-3-3ζ is the 72nd (1680 ppm) out of 19949 proteins. Considered as a whole, the cumulated seven 14-3-3 isoforms even rank within the top 20 (i.e., top 0.1%) most abundant human proteins. However, 14-3-3 isoforms are not uniformly distributed across tissues. Each human cell type has a specific distribution of the 14-3-3 family (Fig. 4H).

We took advantage of the quantified hierarchized affinity profile of 14-3-3 isoforms to build a predictor tool, which estimates the fraction of a given phosphoprotein that is engaged with each distinct 14-3-3 isoform (Supplementary file 1). As an input, the predictor requires (i) the K_D_ of that phosphoprotein for at least one 14-3-3 isoform, and (ii) the cellular concentrations of the seven 14-3-3 isoforms and of the phosphoprotein of interest. The concentrations of a given protein species in a given cell type can be roughly estimated from protein abundance databases (PAXdb database ^3^), by using a simple conversion rule (see Methods).

We used this approach to predict the proportions of each 14-3-3 isoform among the overall 14-3-3/phosphoprotein complexes formed in various tissues, including uterus, rectum and oral cavity, which are all susceptible to hrm-HPV infection, as well as in five other organs (esophagus, skin, lung, brain and heart) (Fig. 4H, I, Supplementary Fig. 8).

Conversely, the free fraction of each phosphoprotein depends on absolute affinity constants. HPV-positive cell lines have been estimated to produce an average of ∼1 ng of E6 per 10^6^ cells, corresponding to an approximate intracellular concentration of 25 nM (^48^ and personal communication from Dr. J. Schweizer, Arborvita Corp, USA). In a situation where the E6 PBM would be fully phosphorylated, we can estimate, using the predictor, that 98%, 85%, 65% or 35% of phosphorylated 33E6, 18E6, 16E6 and 35E6, respectively, would be engaged in 14-3-3 complexes in cells containing the average 14-3-3 concentrations found in human cells (estimated from integrated human data in PAXdb database ^3^) (Supplementary Fig. 8).

## DISCUSSION

E6 oncoproteins of all hrm-HPV types contain a conserved C-terminal PDZ-Binding Motif which can become a potential 14-3-3-binding motif upon phosphorylation ^12, 13, 25^ (Supplementary Fig. 1 and Fig. 1A). Here, we initially set out to analyze the mechanistic and structural basis for the 14-3-3ζ binding to the 18E6 oncoprotein. Comparison to a previously solved complex between 14-3-3σ and HPV16 E6 ^25^ revealed conserved binding principles (Fig. 2B) that are likely to be valid for most hrm-HPV E6/14-3-3 complexes. We also showed that the fusicoccin molecule, a well-known modulator of 14-3-3 interactions, moderately destabilizes E6 binding to 14-3-3 (Fig. 3). This indicated that the hrm-HPV E6/14-3-3 complexes are in principle druggable.

The phosphorylated PBMs of four selected hrm-HPV E6 all detectably bound to 14-3-3 proteins, albeit with surprisingly wide affinity variations spanning a 100-fold K_D_ range for different E6 PBMs binding to a given 14-3-3 isoform (Fig. 1). In the literature, interactions of phosphorylated peptides with 14-3-3 even cover a wider ∼40,000-fold affinity range, from low nanomolar to low millimolar (Fig. 4). As shown in the present work, very modest sequence variations of a phosphopeptide can be sufficient to alter its unbound and/or bound states in a way that greatly impacts binding affinity. Similar principles may govern 14-3-3-binding affinity variations of many other phosphopeptides.

Conversely, the seven isoforms bound each E6 phosphopeptide following a conserved hierarchized profile, with an approximate 11-fold K_D_ ratio between the strongest-binding and the weakest-binding 14-3-3 isoform. Remarkably, 14-3-3s obey the same hierarchy when binding to most of their targets, as supported by our own data on RSK1 and HSPB6 peptides, by our literature curation ^38, 42, 43, 44, 45, 46^, and by the unbiased proteome-wide complexome data very recently made available by the Gygi group in the “Bioplex3” database (https://bioplex.hms.harvard.edu and ^26^) and the human kinome interactome (https://sec-explorer.shinyapps.io/Kinome_interactions/ and ^47^). Only 14-3-3σ may stand out as a partial exception to this rule. While displaying a low affinity to most 14-3-3 targets, it nonetheless binds to a small subset of “proprietary” targets that are not shared with other 14-3-3 isoforms (Fig. 4). This outlier character of 14-3-3σ has already been noticed in previous works dedicated to the structural and functional peculiarities of that isoform ^49, 50^.

We took advantage of the hierarchized target-binding profiles of 14-3-3 isoforms to develop a prediction approach of the 14-3-3 complexome. This approach can compute, for a given cell population, the free and 14-3-3-bound fractions of any phosphoprotein whose cellular concentration and affinity for at least one 14-3-3 isoform are available. The concentration of host proteins can be inferred from the protein abundance databases such as PAXdb (https://pax-db.org and ^3^), while the affinity to a 14-3-3 isoform can easily be obtained using state-of-the-art in vitro protein-peptide binding approaches.

When applied to the rather weakly-expressed HPV E6 proteins, predictions indicated that, in a cellular situation favoring E6 phosphorylation, phospho-E6 molecules should get fully engaged with the 14-3-3 pool for the strongest 14-3-3-binding E6 variants, and only partly engaged for the weaker ones. Such differences are likely to influence the de-phosphorylation kinetics of phospho-E6 molecules from different HPV types, and the subsequent dynamics of cellular mechanisms involving PDZ-containing proteins targeted by E6. We also found that, in tissues susceptible to HPV infections, phosphorylated E6 would be complexed to distinct proportions of 14-3-3 proteins. In particular, phosphorylated E6 might be engaged with a higher proportion of 14-3-3σ in oral cavity, where this otherwise weakly-expressed isoform is particularly abundant.

14-3-3 proteins are abundant in all tissues, yet in variable amounts. It is also known that most tumors adjust their 14-3-3 concentrations, by altering the expression of at least one 14-3-3 isoform ^51, 52, 53^. In all cell types, peaks of bulk phosphorylation occur, for instance at specific cell cycle steps or in reaction to changes in the extracellular environment ^54, 55^. It is tempting to speculate that, as previously proposed by others ^56^, 14-3-3 proteins, notwithstanding their individual functional specificities, may collectively provide a buffering system for intracellular signaling. In such a view, the cumulated concentrations of 14-3-3 are adjusted in each cell type for coping with the most acute phosphorylation outbursts possible in that very cell type. We notice that the highest concentrations of 14-3-3 in human cells are found in platelets (Fig. 4H). Indeed, platelet activation is a phenomenon known to involve powerful phosphorylation events ^57^.

To conclude, the present work opens novel avenues for interpreting, predicting and addressing in a quantitative and global manner the way that distinct 14-3-3 isoforms bind to pools of phosphorylated proteins and thereby modulate their activities.

## METHODS

### Cloning, recombinant protein purification and peptide synthesis

Previously described chimeras contained the C-terminally truncated human 14-3-3σ (Uniprot ID P31947; residues 1-231, 14-3-3σΔC) bearing on its N-terminus a His_6_-tag cleavable by 3C protease and phosphorylatable peptides tethered to the 14-3-3σ C-terminus by a GSGS linker ^33^. The novel chimera was designed taking into account the following modifications. First, it contained the C-terminally truncated human 14-3-3ζ sequence (Uniprot ID P63104; residues 1-229, 14-3-3ζ ΔC) connected to the PKA-phosphorylatable 18E6 heptapeptide around Thr156. Second, the 14-3-3ζ core was modified to block Ser58 phosphorylation (S58A) ^58, 59^. Third, to improve crystallizability, the 14-3-3ζ sequence was mutated by introducing the ^73^EKK^75^→AAA and ^157^KKE^159^→AAA amino acid replacements in the highest-scoring clusters 1 and 2 predicted by the surface entropy reduction approach ^60, 61^. Finally, the linker was changed to GGGG to exclude its unspecific phosphorylation (Supplementary Fig. 3A).

cDNA of the 14-3-3ζ −18E6 chimera was codon-optimized for expression in *Escherichia coli* and synthesized by IDT Technologies (Coralville, Iowa, USA). The 14-3-3ζ ΔC gene was flanked by *NdeI* and *AgeI* restriction endonuclease sites to enable alteration of the 14-3-3 or E6 PBM peptide sequences. The entire 14-3-3ζ −GGGG-18E6 PBM construct was inserted into a pET28-his-3C vector ^62^ using *NdeI* and *XhoI* restriction endonuclease sites. The resulting vector was amplified in DH5α cells and verified using DNA sequencing in Evrogen (Moscow, Russia, www.evrogen.ru).

The assembled vector (Kanamycin resistance) was transformed into chemically competent *E. coli* BL21(DE3) cells for expression either in the absence or in the presence of the His_6_- tagged catalytically active subunit of mouse PKA ^62^. Protein expression was induced by the addition of isopropyl-β-thiogalactoside (IPTG) to a final concentration of 0.5 mM and continued for 16 h at 25 °C. The overexpressed protein was purified using subtractive immobilized metal-affinity chromatography (IMAC) and gel-filtration essentially as described earlier for 14-3-3σ chimeras ^33^ (Supplementary Fig. 3B and C). The purified phosphorylated 14-3-3ζ −18E6 chimera revealed the characteristic downward shift on native PAGE compared to the unphosphorylated counterpart (Supplementary Fig. 3D). Given the absence of PKA phosphorylation sites in the modified 14-3-3ζ core and the linker, this strongly indicated 18E6 phosphorylation by co-expressed PKA. The chimera was fully soluble and stable at concentrations above 20 mg/ml required for crystallization. Protein concentration was determined at 280 nm on a Nanophotometer NP80 (Implen, Germany) using extinction coefficient equal to 0.93 (mg/ml)^-1^ cm^-1^.

For affinity measurements, full-length human 14-3-3 constructs with a rigid N-terminal MBP fusion were used. The coding sequences of the full-length 14-3-3 epsilon, gamma and zeta were received from Prof. Lawrence Banks. cDNAs encoding other full-length 14-3-3 isoforms β, τ, η and σ were obtained as codon-optimized for *E*.*coli* expression synthetic genes from IDT Technologies (Coralville, Iowa, USA). All 14-3-3 isoforms were fused via a three-alanine linker to the C-terminus of a mutant MBP carrying the following amino acid substitutions: D83A, K84A, K240A, E360A, K363A and D364A, as previously described ^63^. All resulting clones were verified by sequencing. The MBP-fused proteins were expressed in *E*.*coli* BL21 with IPTG induction. Proteins were affinity purified on an amylose column and were further purified by ion-exchange chromatography (HiTrap Q HP, GE Healthcare). Protein concentrations were determined by UV spectroscopy. The double-purified samples were supplemented with glycerol and TCEP before aliquoting and freezing in liquid nitrogen.

HPV peptides (35E6: biotin-ttds-SKPTRRETEV; 16E6: biotin-ttds-SSRTRRETQL; 18E6: biotin-ttds-RLQRRRETQV; 33E6: biotin-ttds-SRSRRRETAL; p35E6: biotin-ttds-SKPTRREpTEV; p35E6 E-1A: biotin-ttds-SKPTRREpTAV; p35E6 T-6R: biotin-ttds-SKPRRREpTEV; p35E6 E-1A T-6R: biotin-ttds-SKPRRREpTAV; p16E6: biotin-ttds-SSRTRREpTQL; p18E6: biotin-ttds-RLQRRREpTQV; p33E6: biotin-ttds-SRSRRREpTAL) and RSK1 peptides (RSK1_-1P: biotin-ttds-RRVRKLPSTpTL and RSK1_-2P: biotin-ttds-RRVRKLPSpTTL) were chemically synthesized in-house on an ABI 443A synthesizer with Fmoc strategy. The fluorescently labeled HSPB6 (WLRRApSAPLPGLK) peptide (fpB6) was prepared by FITC labeling of the chemically synthesized peptide as described previously ^25^.

### Fluorescence polarization (FP) assay

Fluorescence polarization was measured with a PHERAstar (BMG Labtech, Offenburg, Germany) microplate reader by using 485 ± 20 nm and 528 ± 20 nm band-pass filters (for excitation and emission, respectively). In direct FP measurements, a dilution series of the 14-3-3 protein was prepared in 96-well plates (96 well skirted pcr plate, 4ti-0740, 4titude, Wotton, UK) in a 20 mM HEPES pH 7.5 buffer containing 150 mM NaCl, 0.5 mM TCEP, 0.01% Tween 20, 50 nM fluorescently-labeled fpB6 peptide and 100 μM fusicoccin (FSC), if indicated. The volume of the dilution series was 40 μl, which was later divided into three technical replicates of 10 μl upon transferring to 384-well micro-plates (low binding microplate, 384 well, E18063G5, Greiner Bio-One, Kremsmünster, Austria). In total, polarization of the probe was measured at 8 different protein concentrations (whereas one contained no protein and corresponded to the free peptide). In competitive FP measurements, the same buffer was supplemented with the protein to achieve a complex formation of 60-80%, based on the titration. Then, this mixture was used for creating a dilution series of the unlabeled competitor (i.e. the studied peptides) and the measurement was carried out identically as in the direct experiment. Analysis of FP experiments were carried out using ProFit, an in-house developed, Python-based fitting program ^64^. The dissociation constant of the direct and competitive FP experiment was obtained by fitting the measured data with quadratic and competitive equation, respectively ^64, 65^. ΔG values were calculated using the formula ΔG=-RT*ln(K_D_), at 295 K. ΔΔG_av_ values were obtained by calculating the average and the standard deviation of all obtained individual ΔΔG values (between different motifs or different proteins), excluding cases when K_D_ > 300 μM.

### Crystallization and structure determination

Crystallization conditions were screened using commercially available and in-house developed kits (Qiagen, Hampton Research, Emerald Biosystems) by the sitting-drop vapor-diffusion method in 96-well MRC 2-drop plates (SWISSCI, Neuheim, Switzerland), using a Mosquito robot (TTP Labtech, Cambridge, UK) at 4 °C. The optimized condition of the crystals consisted of 19% polyethylene glycol 4000, 0.1M cacodylate buffered at pH 5.5. For soaking, crystals were transferred to a mother-liquor solution containing (saturated, partially precipitated) 5 mM fusicoccin and crystals were harvested after an 18h incubation period. All crystals were flash-cooled in a cryoprotectant solution containing 20% glycerol and stored in liquid nitrogen.

X-ray diffraction data were collected at the Synchrotron Swiss Light Source (SLS) (Switzerland) on the X06DA (PXIII) beamline and processed with the program XDS ^66^. The crystal structure was solved by molecular replacement with a high-resolution crystal structure of 14-3-3ζ (PDB ID 2O02) using Phaser ^67^ and structure refinement was carried out with PHENIX ^68^. TLS refinement was applied during the refinement. The crystallographic parameters and the statistics of data collection and refinement are shown in Table 1.

### Predictions of proportions of 14-3-3 isoform complexes

We built a simple predictor tool that can be run using the Excel program (Supplementary file 1). As an input, the predictor requires (i) the K_D_ of binding of the phosphoprotein of interest for at least one 14-3-3 isoform, and (ii) the cellular concentrations of the seven 14-3-3 isoforms and of the phosphoprotein of interest. As an output, the predictor estimates the fraction of phosphoprotein that is engaged with each distinct 14-3-3 isoform.

From the provided K_D_ value(s), the predictor derives all K_D_ values for the remaining 14-3-3 isoforms, using the average relative affinity ratios described in our results.

The cellular protein concentrations required by the predictor can either be determined experimentally or fixed arbitrarily to explore hypotheses. In the present work, we used “integrated” protein abundance data from the PAXdb database ^3^. In this database, abundance of a given protein is expressed as the ppm fraction of the number of molecules of that protein species relative to the cumulated number of all molecules of all protein species detected in the sample ^69^. For instance, if the abundance of a protein species Prot_n_ is Ab_Protn_ = 1000 ppm, this means that over a total one million (10^6^) counted proteins, one thousand (10^3^) correspond to the protein species of interest. Furthermore, the total intracellular protein concentration Prot_Tot_ has been estimated to be around 3 mM (https://bionumbers.hms.harvard.edu/keynumbers.aspx). Therefore, for any protein Prot_n_ of interest, one can use the ppm abundance value, Ab_Protn_, to roughly estimate the cellular molar concentration of that protein (Prot_n_) using the following formula:

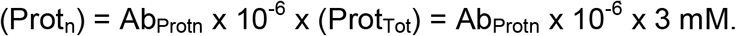

For instance,

for Ab_Protn_ = 1 ppm, (Prot_n_) = 1 × 10^−6^ x 3 mM = 3 nM

for Ab_Protn_ = 1000 ppm, (Prot_n_) = 1000 × 10^−6^ x 3 mM = 3 µM.

While 3 mM is a reasonable estimate of the total intracellular protein concentration, one might argue that picking this particular value is an arbitrary choice, since total numbers of protein molecules may vary by up to 10-fold from one cell type to another (http://book.bionumbers.org/how-many-proteins-are-in-a-cell/). However, in practice, we found that the proportions of bound 14-3-3 isoforms computed by the predictor do not significantly change for any value of (Prot_Tot_) taken in the range between 1 mM and 10 mM.

## Supporting information

Predictor of the 14-3-3 complexomes

## Acknowledgments

We would like to thank Prof. Lawrence Banks for the shared plasmids, Prof. Alexey Babakov for the provided fusicoccin preparations and Dr. Yaroslav Faletrov for fusicoccin identity verification. N.N.S. is grateful to the Russian Science Foundation for the grant no. 19-74-10031. We thank the support of the Swiss Light Source synchrotron (P. Scherrer Institute, Villigen, Switzerland) and the help of the beam-scientist at the PXIII beamline. The work was supported by the Ligue contre le cancer (équipe labellisée 2015 to G.T.), the French Infrastructure for Integrated Structural Biology (FRISBI), and Instruct-ERIC. G.G. was supported by the Post-doctorants en France program of the Fondation ARC.

## Author contributions

G.G. purified proteins, carried out FP experiments, performed crystallographic and bioinformatics studies, analyzed the data and edited the paper. K.V.T. cloned, purified and characterized proteins. C.K. cloned and purified proteins. P.E. synthesized the peptides. G.T. performed data analysis, data interpretation, wrote and edited the paper. N.N.S. contributed to protein purification and crystallographic experiments, supervised the research, analyzed the data, wrote and edited the paper.

## Data availability

The refined model and the structure factor amplitudes have been deposited in the PDB with the accession codes 6ZFD and 6ZFG. All other data supporting the findings of this study are available from the corresponding authors upon reasonable request.

## References

1. Aitken A. 14-3-3 proteins: a historic overview. Semin Canc Biol 16, 162–172 (2006).

2. Boston PF, Jackson P, Thompson RJ. Human 14-3-3 protein: radioimmunoassay, tissue distribution, and cerebrospinal fluid levels in patients with neurological disorders. J Neurochem 38, 1475–1482 (1982).

3. Wang M, Herrmann CJ, Simonovic M, Szklarczyk D, von Mering C. Version 4.0 of PaxDb: Protein abundance data, integrated across model organisms, tissues, and cell-lines. Proteomics 15, 3163–3168 (2015).

4. Muslin AJ, Tanner JW, Allen PM, Shaw AS. Interaction of 14-3-3 with signaling proteins is mediated by the recognition of phosphoserine. Cell 84, 889–897 (1996).

5. Yaffe MB, et al. The structural basis for 14-3-3:phosphopeptide binding specificity. Cell 91, 961–971 (1997).

6. Ganguly S, Weller J, Ho A, Chemineau P, Malpaux B, Klein D. Melatonin synthesis: 14-3-3-dependent activation and inhibition of arylalkylamine N-acetyltransferase mediated by phosphoserine-205. Proc Natl Acad Sci U S A 102, 1222–1227 (2005).

7. Paiardini A, et al. The phytotoxin fusicoccin differently regulates 14-3-3 proteins association to mode III targets. IUBMB Life 66, 52–62 (2014).

8. Obsil T, Obsilova V. Structural basis of 14-3-3 protein functions. Semin Cell Dev Biol 22, 663–672 (2011).

9. Mackintosh C. Dynamic interactions between 14-3-3 proteins and phosphoproteins regulate diverse cellular processes. Biochem J 381, 329–342 (2004).

10. Stevers LM, et al. Modulators of 14-3-3 Protein-Protein Interactions. J Med Chem 61, 3755–3778 (2018).

11. Nathan KG, Lal SK. The Multifarious Role of 14-3-3 Family of Proteins in Viral Replication. Viruses 12, (2020).

12. Boon SS, Banks L. High-risk human papillomavirus E6 oncoproteins interact with 14-3-3zeta in a PDZ binding motif-dependent manner. J Virol 87, 1586–1595 (2013).

13. Boon SS, Tomaic V, Thomas M, Roberts S, Banks L. Cancer-causing human papillomavirus E6 proteins display major differences in the phospho-regulation of their PDZ interactions. J Virol 89, 1579–1586 (2015).

14. Ganti K, et al. The Human Papillomavirus E6 PDZ Binding Motif: From Life Cycle to Malignancy. Viruses 7, 3530–3551 (2015).

15. Basukala O, Sarabia-Vega V, Banks L. Human papillomavirus oncoproteins and post-translational modifications: generating multifunctional hubs for overriding cellular homeostasis. Biological chemistry 401, 585–599 (2020).

16. McBride AA. Oncogenic human papillomaviruses. Philos Trans R Soc Lond B Biol Sci 372, (2017).

17. Vande Pol SB, Klingelhutz AJ. Papillomavirus E6 oncoproteins. Virology 445, 115–137 (2013).

18. Suarez I, Trave G. Structural Insights in Multifunctional Papillomavirus Oncoproteins. Viruses 10, (2018).

19. Poirson J, et al. Mapping the interactome of HPV E6 and E7 oncoproteins with the ubiquitin-proteasome system. FEBS J 284, 3171–3201 (2017).

20. Celegato M, et al. A novel small-molecule inhibitor of the human papillomavirus E6-p53 interaction that reactivates p53 function and blocks cancer cells growth. Cancer Lett 470, 115–125 (2020).

21. Kolluru S, Momoh R, Lin L, Mallareddy JR, Krstenansky JL. Identification of potential binding pocket on viral oncoprotein HPV16 E6: a promising anti-cancer target for small molecule drug discovery. BMC molecular and cell biology 20, 30 (2019).

22. Zanier K, et al. The E6AP binding pocket of the HPV16 E6 oncoprotein provides a docking site for a small inhibitory peptide unrelated to E6AP, indicating druggability of E6. PLoS One 9, e112514 (2014).

23. Ramirez J, et al. Targeting the Two Oncogenic Functional Sites of the HPV E6 Oncoprotein with a High-Affinity Bivalent Ligand. Angew Chem Int Ed Engl 54, 7958–7962 (2015).

24. Songyang Z, et al. Recognition of unique carboxyl-terminal motifs by distinct PDZ domains. Science 275, 73–77 (1997).

25. Gogl G, et al. Dual Specificity PDZ- and 14-3-3-Binding Motifs: A Structural and Interactomics Study. Structure 28, 747–759 e743 (2020).

26. Huttlin EL, et al. Dual Proteome-scale Networks Reveal Cell-specific Remodeling of the Human Interactome. bioRxiv, 2020.2001.2019.905109 (2020).

27. Luck K, Charbonnier S, Trave G. The emerging contribution of sequence context to the specificity of protein interactions mediated by PDZ domains. FEBS Lett 586, 2648–2661 (2012).

28. Puntervoll P, et al. ELM server: A new resource for investigating short functional sites in modular eukaryotic proteins. Nucleic Acids Res 31, 3625–3630 (2003).

29. Kuhne C, Gardiol D, Guarnaccia C, Amenitsch H, Banks L. Differential regulation of human papillomavirus E6 by protein kinase A: conditional degradation of human discs large protein by oncogenic E6. Oncogene 19, 5884–5891 (2000).

30. Miller CJ, Turk BE. Homing in: Mechanisms of Substrate Targeting by Protein Kinases. Trends Biochem Sci 43, 380–394 (2018).

31. Ben-Shimon A, Niv MY. Deciphering the Arginine-binding preferences at the substrate-binding groove of Ser/Thr kinases by computational surface mapping. PLoS Comput Biol 7, e1002288 (2011).

32. Espejo AB, et al. PRMT5 C-terminal Phosphorylation Modulates a 14-3-3/PDZ Interaction Switch. J Biol Chem 292, 2255–2265 (2017).

33. Sluchanko NN, Tugaeva KV, Greive SJ, Antson AA. Chimeric 14-3-3 proteins for unraveling interactions with intrinsically disordered partners. Sci Rep 7, 12014 (2017).

34. Tugaeva KV, Remeeva A, Gushchin I, Cooley RB, Sluchanko NN. Design, expression, purification and crystallization of human 14-3-3zeta protein chimera with phosphopeptide from proapoptotic protein BAD. Protein Expr Purif, 105707 (2020).

35. Gogl G, et al. Dynamic control of RSK complexes by phosphoswitch-based regulation. FEBS J 285, 46–71 (2018).

36. Camoni L, Visconti S, Aducci P. The phytotoxin fusicoccin, a selective stabilizer of 14-3-3 interactions? IUBMB Life 65, 513–517 (2013).

37. De Vries-van Leeuwen IJ, et al. Interaction of 14-3-3 proteins with the estrogen receptor alpha F domain provides a drug target interface. Proc Natl Acad Sci U S A 110, 8894–8899 (2013).

38. Stevers LM, et al. Characterization and small-molecule stabilization of the multisite tandem binding between 14-3-3 and the R domain of CFTR. Proc Natl Acad Sci U S A 113, E1152–1161 (2016).

39. Wurtele M, Jelich-Ottmann C, Wittinghofer A, Oecking C. Structural view of a fungal toxin acting on a 14-3-3 regulatory complex. EMBO J 22, 987–994 (2003).

40. Sengupta A, Liriano J, Miller BG, Frederich JH. Analysis of Interactions Stabilized by Fusicoccin A Reveals an Expanded Suite of Potential 14-3-3 Binding Partners. ACS Chem Biol 15, 305–310 (2020).

41. Kaplan A, et al. Polypharmacological Perturbation of the 14-3-3 Adaptor Protein Interactome Stimulates Neurite Outgrowth. Cell chemical biology 27, 657–667 e656 (2020).

42. Kilisch M, Lytovchenko O, Arakel EC, Bertinetti D, Schwappach B. A dual phosphorylation switch controls 14-3-3-dependent cell surface expression of TASK-1. J Cell Sci 129, 831–842 (2016).

43. Centorrino F, Ballone A, Wolter M, Ottmann C. Biophysical and structural insight into the USP8/14-3-3 interaction. FEBS Lett 592, 1211–1220 (2018).

44. Manschwetus JT, et al. Binding of the Human 14-3-3 Isoforms to Distinct Sites in the Leucine-Rich Repeat Kinase 2. Front Neurosci 14, 302 (2020).

45. Wolter M, et al. Selectivity via Cooperativity: Preferential Stabilization of the p65/14-3-3 Interaction with Semisynthetic Natural Products. J Am Chem Soc 142, 11772–11783 (2020).

46. Rose R, Rose M, Ottmann C. Identification and structural characterization of two 14-3-3 binding sites in the human peptidylarginine deiminase type VI. J Struct Biol 180, 65–72 (2012).

47. Buljan M, et al. Kinase Interaction Network Expands Functional and Disease Roles of Human Kinases. Molecular Cell 79, 504-520.e509 (2020).

48. Peck RB, et al. A Magnetic Immunochromatographic Strip Test for Detection of Human Papillomavirus 16 E6. Clinical Chemistry 52, 2170–2172 (2006).

49. Wilker E, Grant R, Artim S, Yaffe M. A structural basis for 14-3-3sigma functional specificity. J Biol Chem 280, 18891–18898 (2005).

50. Benzinger A, Popowicz G, Joy J, Majumdar S, Holak T, Hermeking H. The crystal structure of the non-liganded 14-3-3sigma protein: insights into determinants of isoform specific ligand binding and dimerization. Cell research 15, 219–227 (2005).

51. Uchida D, Begum NM, Almofti A, Kawamata H, Yoshida H, Sato M. Frequent downregulation of 14-3-3 sigma protein and hypermethylation of 14-3-3 sigma gene in salivary gland adenoid cystic carcinoma. Br J Cancer 91, 1131–1138 (2004).

52. Nakanishi K, Hashizume S, Kato M, Honjoh T, Setoguchi Y, Yasumoto K. Elevated expression levels of the 14-3-3 family of proteins in lung cancer tissues. Hum Antibodies 8, 189–194 (1997).

53. Nishimura Y, et al. Overexpression of YWHAZ relates to tumor cell proliferation and malignant outcome of gastric carcinoma. Br J Cancer 108, 1324–1331 (2013).

54. Benz C, Urbaniak MD. Organising the cell cycle in the absence of transcriptional control: Dynamic phosphorylation co-ordinates the Trypanosoma brucei cell cycle post-transcriptionally. PLoS Pathog 15, e1008129 (2019).

55. Carpy A, et al. Absolute proteome and phosphoproteome dynamics during the cell cycle of Schizosaccharomyces pombe (Fission Yeast). Mol Cell Proteomics 13, 1925–1936 (2014).

56. Grant MP, Cavanaugh A, Breitwieser GE. 14-3-3 Proteins Buffer Intracellular Calcium Sensing Receptors to Constrain Signaling. PLoS One 10, e0136702 (2015).

57. Babur O, et al. Phosphoproteomic quantitation and causal analysis reveal pathways in GPVI/ITAM-mediated platelet activation programs. Blood, (2020).

58. Gu Y-M, Jin Y-H, Choi J-K, Baek K-H, Yeo C-Y, Lee K-Y. Protein kinase A phosphorylates and regulates dimerization of 14-3-3 epsilon. FEBS Lett 580, 305–310 (2006).

59. Sluchanko NN, Uversky VN. Hidden disorder propensity of the N-terminal segment of universal adapter protein 14-3-3 is manifested in its monomeric form: Novel insights into protein dimerization and multifunctionality. Biochim Biophys Acta 1854, 492–504 (2015).

60. Goldschmidt L, Cooper DR, Derewenda ZS, Eisenberg D. Toward rational protein crystallization: A Web server for the design of crystallizable protein variants. Protein Sci 16, 1569–1576 (2007).

61. Goldschmidt L, Cooper DR, Derewenda ZS, Eisenberg D. SERp Server. (ed^(eds) (2007).

62. Tugaeva KV, Tsvetkov PO, Sluchanko NN. Bacterial co-expression of human Tau protein with protein kinase A and 14-3-3 for studies of 14-3-3/phospho-Tau interaction. PLoS One 12, e0178933 (2017).

63. Zanier K, et al. Structural basis for hijacking of cellular LxxLL motifs by papillomavirus E6 oncoproteins. Science 339, 694–698 (2013).

64. Simon MA, et al. High-throughput competitive fluorescence polarization assay reveals functional redundancy in the S100 protein family. FEBS J 287, 2834–2846 (2020).

65. Roehrl MH, Wang JY, Wagner G. A general framework for development and data analysis of competitive high-throughput screens for small-molecule inhibitors of protein-protein interactions by fluorescence polarization. Biochemistry 43, 16056–16066 (2004).

66. Kabsch W. Xds. Acta Crystallogr D Biol Crystallogr 66, 125–132 (2010).

67. McCoy AJ, Grosse-Kunstleve RW, Adams PD, Winn MD, Storoni LC, Read RJ. Phaser crystallographic software. J Appl Crystallogr 40, 658–674 (2007).

68. Adams PD, et al. PHENIX: a comprehensive Python-based system for macromolecular structure solution. Acta Crystallogr D Biol Crystallogr 66, 213–221 (2010).

69. Wang M, et al. PaxDb, a database of protein abundance averages across all three domains of life. Mol Cell Proteomics 11, 492–500 (2012).

