## Supplementary Table 1, Supplementary Figs 1-10 for "Hierarchized phosphotarget binding by the seven human 14-3-3 isoforms"

### SUPPLEMENTARY INFORMATION

*Gogl et al.*

**Supplementary Table 1.** Crystal structures of the 14-3-3/motif III peptide complexes available in the PDB. FSC – fusicoccin, cotA – cotylenin A, “modified” indicates that the peptide sequence or length are not authentic. Red bold font highlights the peptides matching the pS/pT**XX**-COOH motif III consensus.

| PDB ID | Motif III peptide | (pS/pT)Xn-COOH, n= | 14-3-3 partner | Complex contents | Resolution | Ref. |
|---|---|---|---|---|---|---|
| **1O9D** | ..QSY**pT**V-COOH | 1 | PMA2 | 1433 + peptide | **2.30 Å** | 1 |
| **1O9F** | ..QSY**pT**V-COOH | 1 | PMA2 | 1433 + peptide +FSC | **2.70 Å** | 1 |
| **3E6Y** | ..QSY**pT**V-COOH | 1 | PMA2 | 1433 + peptide +cotA | **2.50 Å** | 2 |
| **5NWI** | ..YFS**pS**N-COOH | 1 | KAT1 | 1433 + peptide | **2.35 Å** | 3 |
| **5NWJ** | ..YFS**pS**N-COOH | 1 | KAT1 | 1433 + peptide | **2.07 Å** | 3 |
| **5NWK** | ..YFS**pS**N-COOH | 1 | KAT1 | 1433 + peptide +FSC | **3.30 Å** | 3 |
| **3IQU** | .QRST**pS**T-COOH | 1 | Raf1 (modified) | 1433 + peptide | **1.05 Å** | 4 |
| **3IQV** | .QRST**pS**T-COOH | 1 | Raf1 (modified) | 1433 + peptide +FSC | **1.20 Å** | 4 |
| **3P1N** | .KRRK**pS**V-COOH | 1 | TASK-3 | 1433 + peptide | **1.40 Å** | 5 |
| **3P1O** | .KRRK**pS**V-COOH | 1 | TASK-3 | 1433 + peptide +FSC | **1.90 Å** | 5 |
| **3P1P** | .KRRK**pS**V-COOH | 1 | TASK-3 | 1433 mutant + peptide | **1.95 Å** | 5 |
| **3P1Q** | .KRRK**pS**V-COOH | 1 | TASK-3 | 1433 mutant + peptide +FSC | **1.70 Å** | 5 |
| **3P1R** | .KRRK**pS**V-COOH | 1 | TASK-3 | 1433 mutant + peptide | **1.70 Å** | 5 |
| **3P1S** | .KRRK**pS**V-COOH | 1 | TASK-3 | 1433 mutant + peptide +FSC | **1.65 Å** | 5 |
| **3SMK** | .KRRK**pS**V-COOH | 1 | TASK-3 | 1433 mutant + peptide +cotA | **2.10 Å** | 5 |
| **3SML** | .KRRK**pS**V-COOH | 1 | TASK-3 | 1433 mutant + peptide +FSC_deriv. | **1.90 Å** | 5 |
| **3SMN** | .KRRK**pS**V-COOH | 1 | TASK-3 | 1433 mutant + peptide +FSC_deriv. | **2.00 Å** | 5 |
| **3SP5** | .KRRK**pS**V-COOH | 1 | TASK-3 | 1433 mutant + peptide +cotA_deriv. | **1.80 Å** | 5 |
| **3SPR** | .KRRK**pS**V-COOH | 1 | TASK-3 | 1433 mutant + peptide +FSC_deriv. | **1.99 Å** | 5 |
| **3UX0** | .KRRK**pS**V-COOH | 1 | TASK-3 | 1433 mutant + peptide +FSC_deriv. | **1.75 Å** | 5 |
| **6GHP** | .KRRK**pS**V-COOH | 1 | TASK-3 | 1433 mutant + FSC_deriv. | **1.95 Å** | 6 |
| **3SMM** | .KRRK**pS**V-COOH | 1 | TASK-3 | 1433 mutant + FSC_deriv. | **2.00 Å** | 5 |
| **3SMO** | .KRRK**pS**V-COOH | 1 | TASK-3 | 1433 mutant + FSC_deriv. | **1.80 Å** | 5 |
| **4FR3** | .KRRK**pS**V-COOH | 1 | TASK-3 | 1433 mutant + FSC_deriv. | **1.90 Å** | 5 |
| **4JC3** | .GFPA**pT**V-COOH | 1 | ERα | 1433 + peptide | **2.05 Å** | 7 |
| **4JDD** | .GFPA**pT**V-COOH | 1 | ERα | 1433 + peptide +FSC | **2.10 Å** | 7 |
| **5N10** | .GFPA**pT**V-COOH | 1 | ERα | 1433 + peptide +ring stabilizer | **1.60 Å** | 8 |
| **6HHP** | .GFPA**pT**V-COOH | 1 | ERα | 1433 + peptide +stabilizer 1 | **1.80 Å** | 9 |
| **6HMT** | .GFPA**pT**V-COOH | 1 | ERα | 1433 + peptide +stabilizer 2 | **1.10 Å** | 9 |
| **6HKB** | .GFPA**pT**V-COOH | 1 | ERα | 1433 + peptide +stabilizer 3 | **1.70 Å** | 9 |
| **6HKF** | .GFPA**pT**V-COOH | 1 | ERα | 1433 + peptide +stabilizer 4 | **1.80 Å** | 9 |
| **6HN2** | .GFPA**pT**V-COOH | 1 | ERα | 1433 + peptide +stabilizer 5 | **1.70 Å** | 9 |
| **6HMU** | .GFPA**pT**V-COOH | 1 | ERα | 1433 + peptide +stabilizer 6 | **1.20 Å** | 9 |
| **6W0L** | SMRRM**pS**M-COOH | 1 | Henipah virus protein W | 1433 + peptide | **2.30 Å** | 10 |
| **4O46** | **RTARpSKV-COOH** | **2** | **Influenza virus NS1** | **1433 + peptide** | **2.90 Å** | 11 |
| **6TWZ** | **RTRREpTQL-COOH** | **2** | **HPV16 E6** | **1433 + peptide** | **2.80 Å** | 12 |
| **6T80** | LRRN**pS**GCG-COOH | 3 | AANAT (modified) | 1433σ-peptide chimera | **2.99 Å** | 13 |

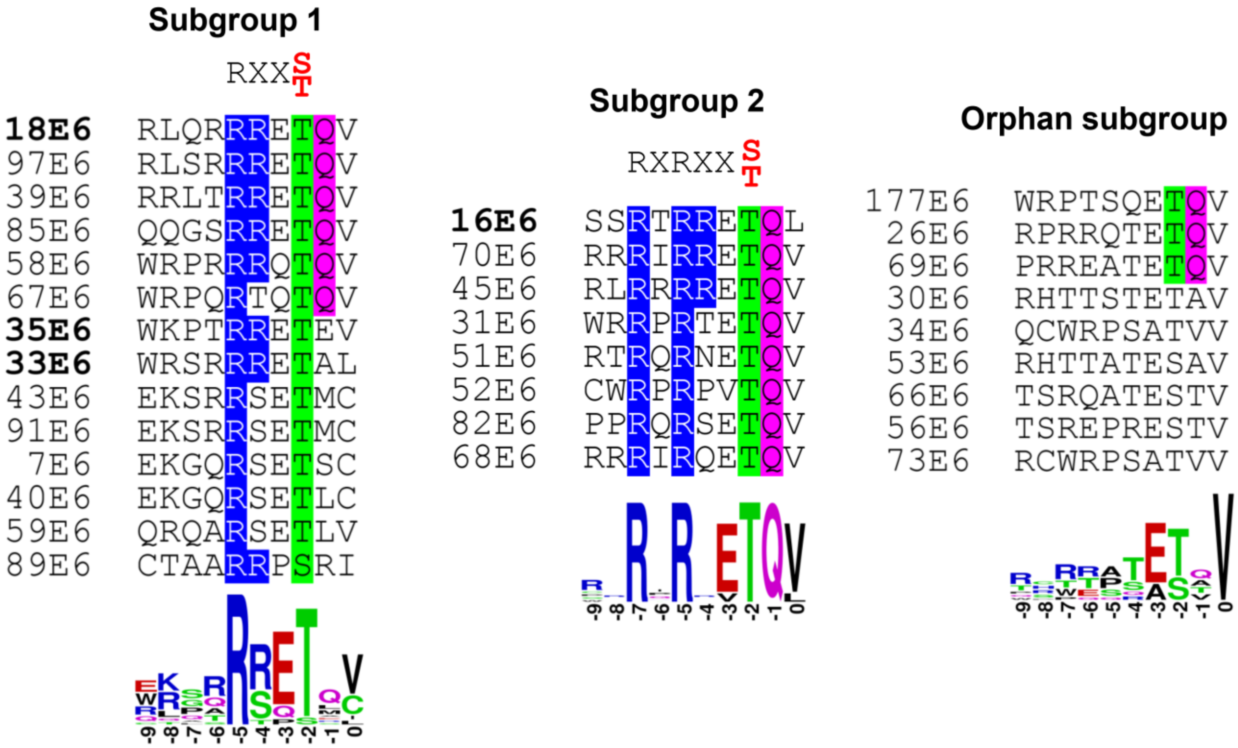


**Supplementary figure 1**. Classification of 31 PBM-containing HPV-E6 proteins based on the correspondence of their C-terminal PBMs to the consensus motifs phosphorylatable by basophilic kinases (subgroups 1 and 2). The third, orphan subgroup comprises PBMs of HPV-E6 proteins whose phosphorylation is less certain. Bold font marks the E6 PBMs used in this work. Note that many PBMs overlap with recognition motifs for phosphorylation by DNA damage response kinases ATM/ATR (TQ and SQ sites highlighted). Below are shown Weblogo diagrams [14] for the PBMs within each of the three subgroups (positions are numbered according to the PBM convention).

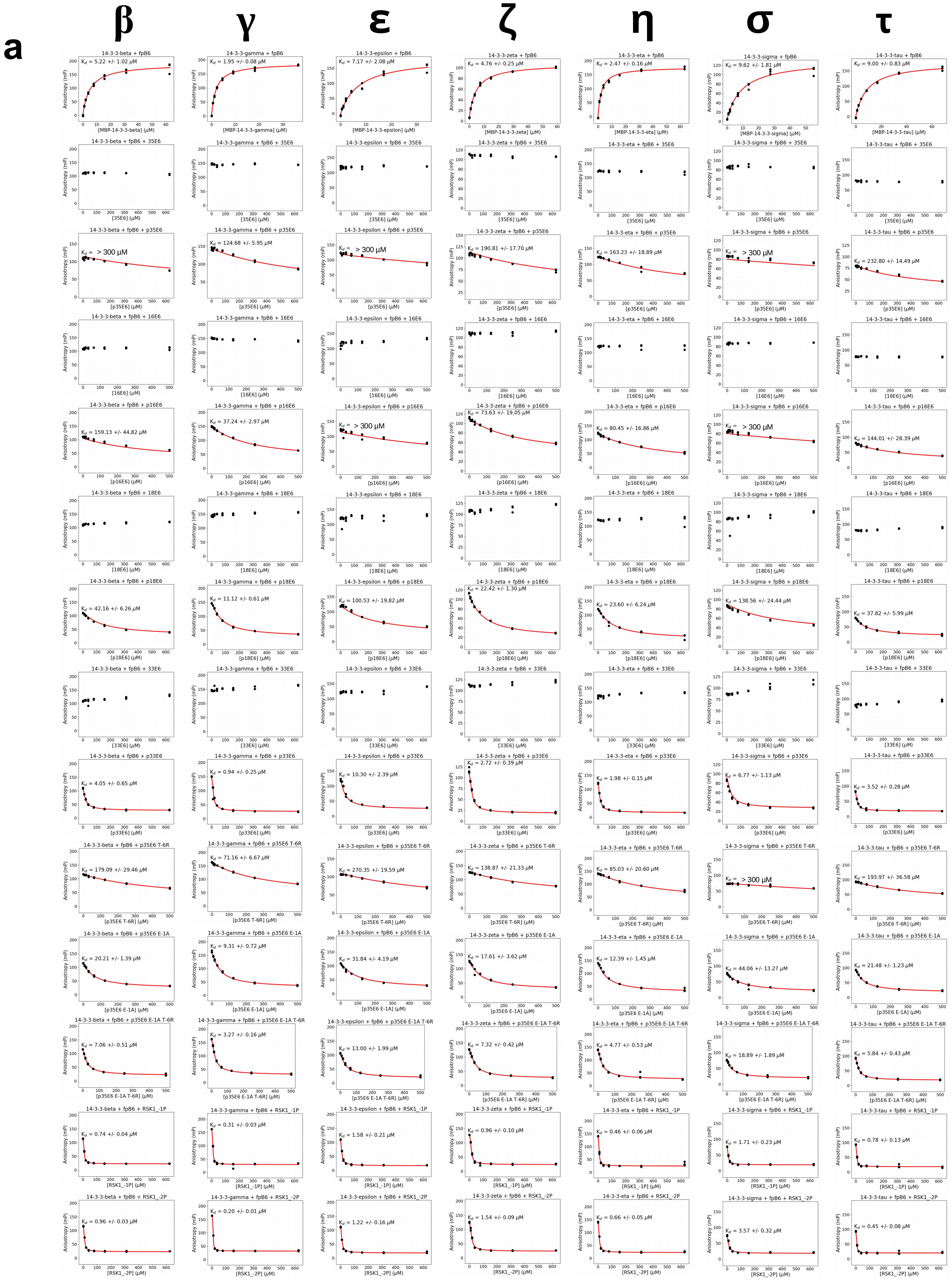


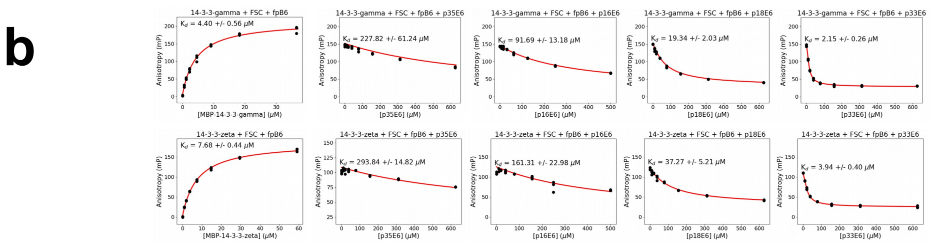


**Supplementary Fig. 2**
See the legend on the next page

**Supplementary figure 2.** Competitive FP measurements with the entire family of seven human full-length 14-3-3 proteins. **A**. In each section, the first panel shows the direct FP experiments between the labeled peptide tracer and the titrated 14-3-3 protein and the following panels show competitive titrations. Competitive experiments were performed at a relatively high protein concentration to achieve 80% complex formation with the peptide tracer. Obtained polarization values were fitted with ProFit [15]. During competitive fitting, an experimental window close to the window of the direct experiment was either achieved without restraints or a restrained fitting was used. No fitted curve is shown if we did not observe a quantifiable competition. **B**. Direct and competitive FP experiments in the presence of FSC.

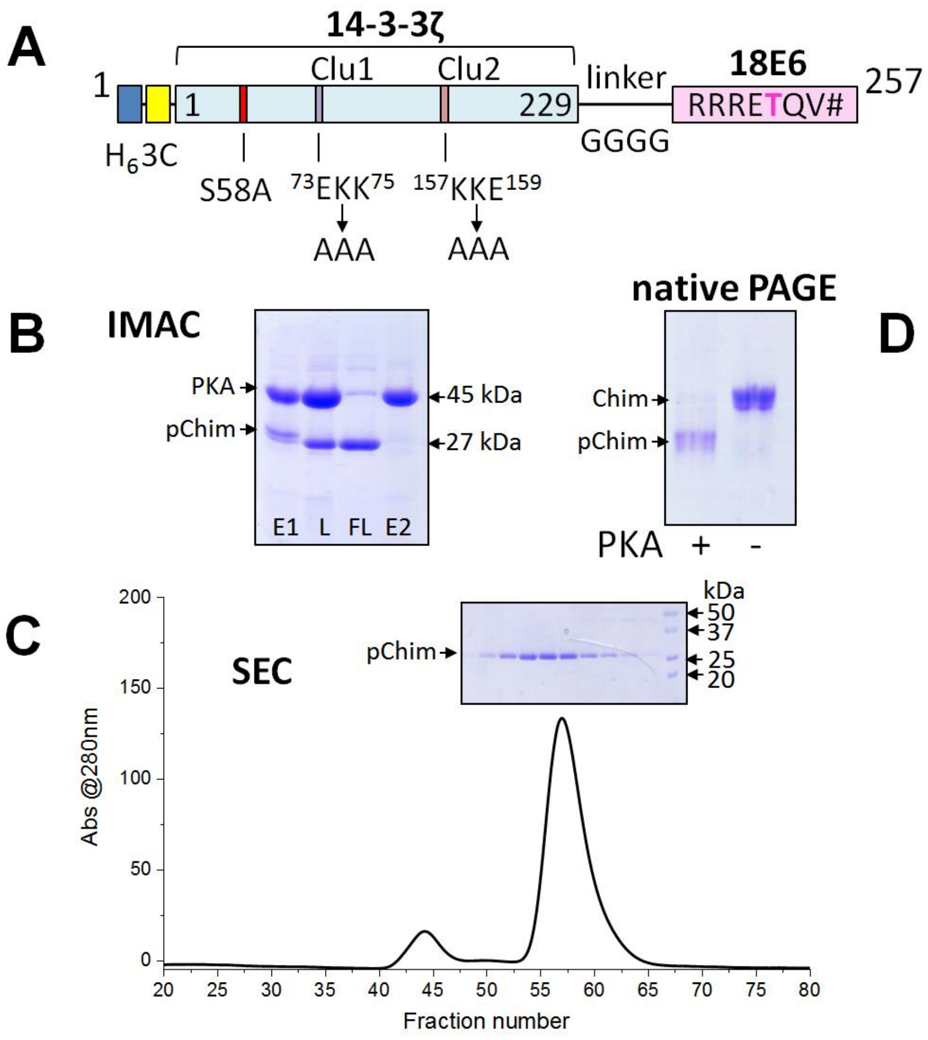


**Supplementary figure 3**. Design and purification of the 14-3-3ζ chimera with the 18E6 phosphopeptide. **A**. Schematic representation of the primary structure of the chimera. The His-tag (H6), 3C protease cleavage site (3C), 14-3-3ζ core modified to prevent phosphorylation of Ser58 and to promote crystallization by the surface entropy reducing mutations (highest scoring clusters 1 and 2 (clu1 and clu2, respectively) are marked [16, 17]), the GGGG linker and the 18E6 C-terminal phosphorylatable peptide are indicated (position of the phosphorylatable threonine is highlighted). The C-terminal carboxylic group is denoted by # and corresponds to the C-terminus of the 18E6 protein. **B**. Purification of the chimera bacterially co-expressed with the His-tagged catalytically active mouse PKA subunit by subtractive immobilized metal affinity chromatography (IMAC) analyzed by SDS-PAGE: E1 – fraction bound to the HisTrap HP column, L – E1 fraction proteolyzed by 3C and loaded on the HisTrap HP column again, FL – unbound fraction containing the untagged phosphorylated chimera, E2 – His-tag containing proteins bound to the column again. Positions of PKA and chimera as well as their apparent Mw values are shown by arrows. Note the downward shift of the chimera band due to 3C treatment. **C**. Size-exclusion chromatography (SEC) profile of the chimera on a Superdex 75 26/60 column (GE Healthcare) shown with the analysis of protein content of the main peak by SDS-PAGE. Arrows indicate positions of the chimera and Mw markers (shown in kDa). **D**. Native PAGE analysis of the purified chimera expressed in *E. coli* in the presence (+) or in the absence of PKA (-). Note the higher electrophoretic mobility of the phosphorylated chimera due to additional negative charges conferred by phosphate moiety.

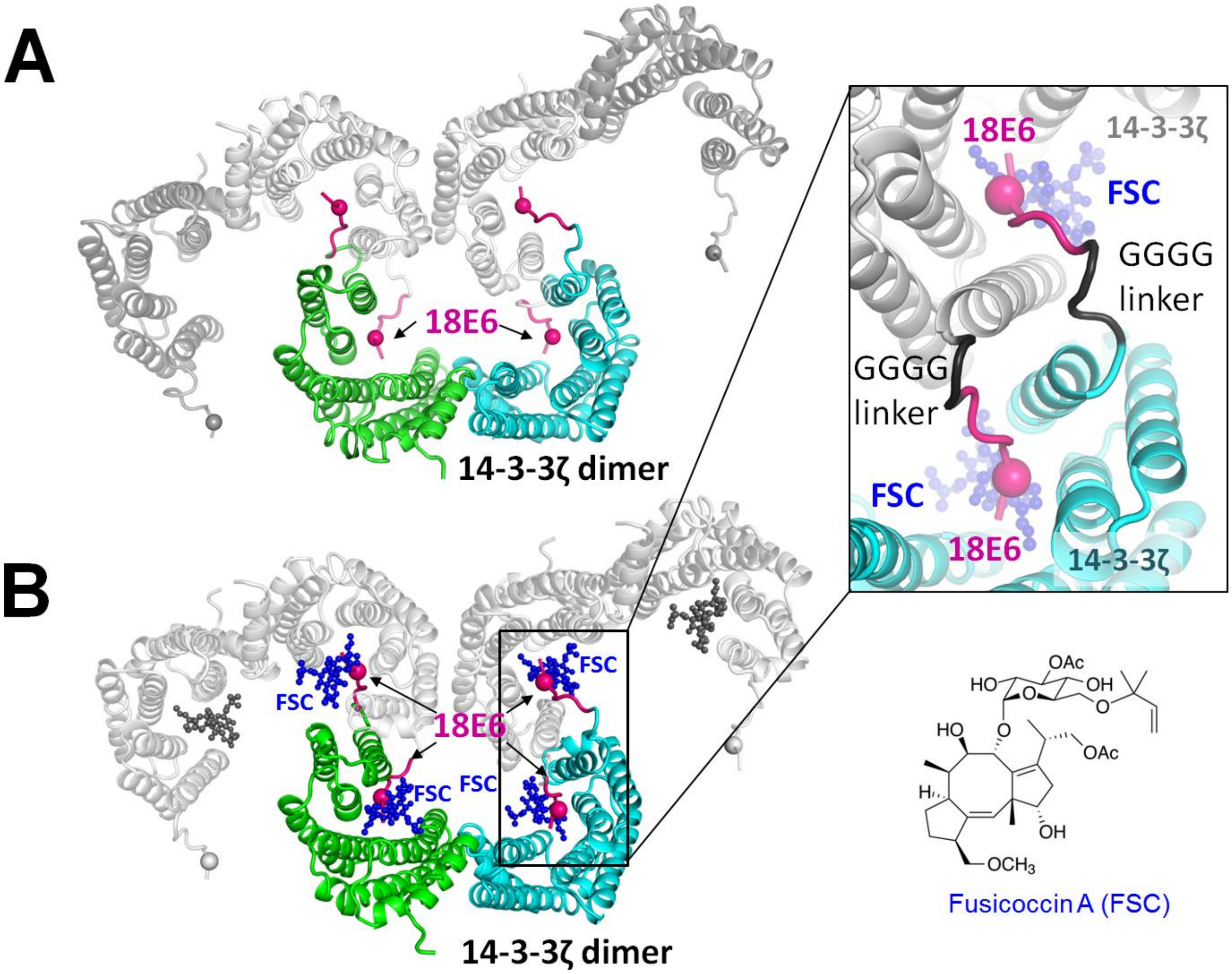


**Supplementary figure 4**. The arrangement of the chimera molecules in the crystal structures obtained in the absence (**A**) or in the presence of FSC (**B**). One 14-3-3ζ dimer of the asymmetric unit is colored, its crystallographically symmetric dimers are light grey. 18E6 phosphopeptides (magenta) are indicated by arrows, phospho-Thr residues are shown by spheres. The interdimer phosphopeptide swap stabilizing the supramolecular assembly is shown in a magnified view only for the FSC bound structure. A chemical formula of FSC is shown in the bottom-right corner.

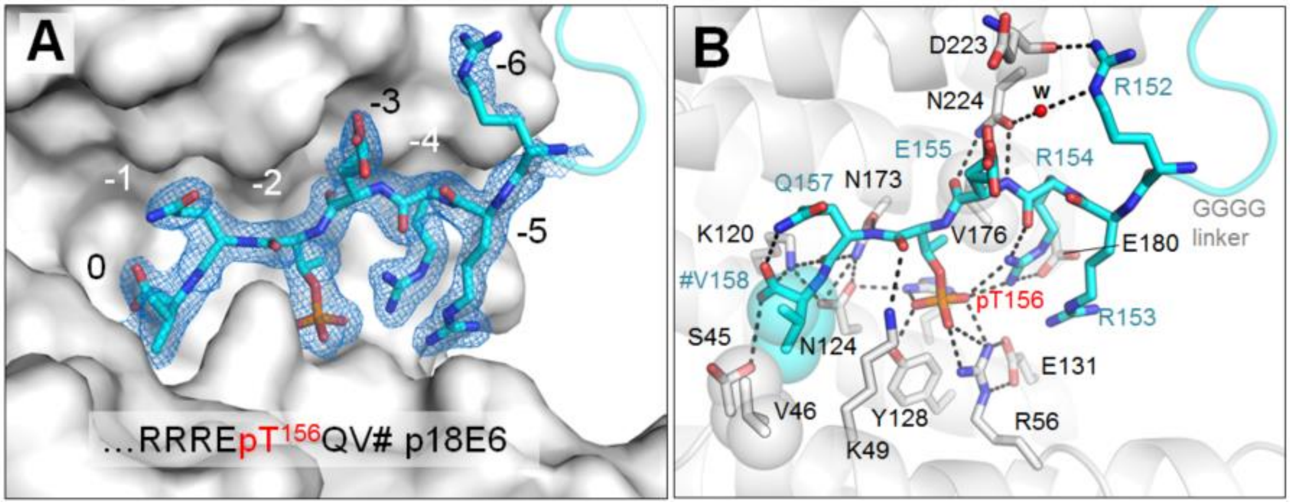


**Supplementary figure 5**. Molecular interface between 14-3-3ζ and phosphorylated 18E6 PBM at a 1.9 Å resolution. **A**. A magnified view on one of the amphipathic grooves of 14-3-3ζ showing the conformation of the 18E6 phosphopeptide and the corresponding $2F_o$-$F_c$ electron density maps contoured at 1σ. Positions are numbered according to the PBM convention. **B**. Polar contacts (dashed lines) and hydrophobic interactions (semitransparent spheres) stabilizing the bound 18E6 peptide conformation.

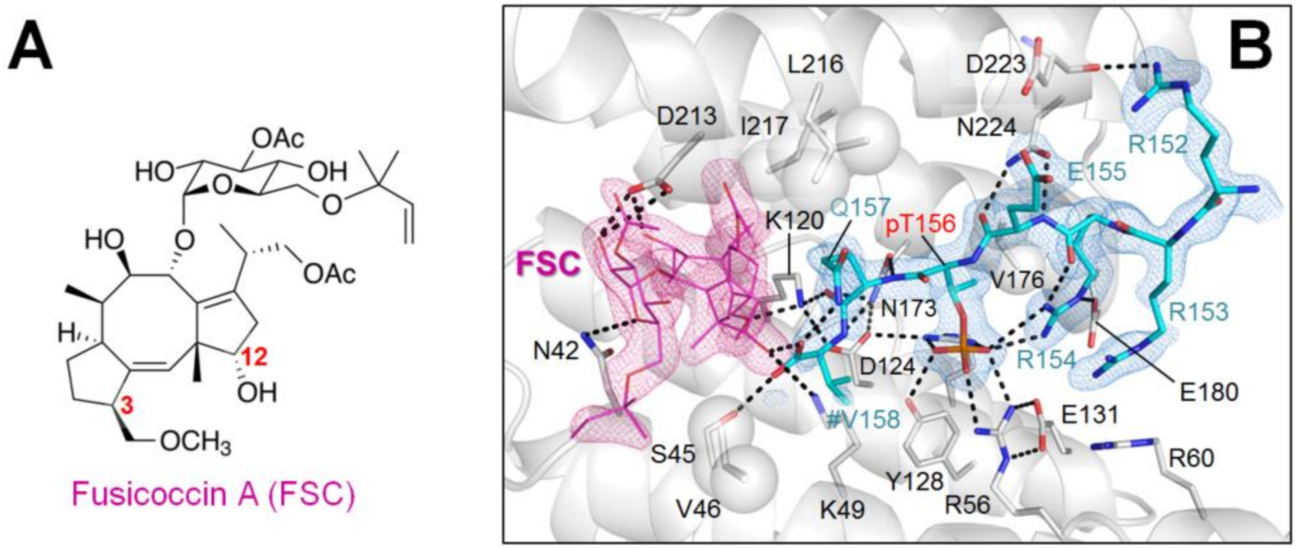


**Supplementary figure 6**. Crystal structure of the ternary complex 14-3-3ζ/18E6 PBM/FSC. A. Chemical formula of FSC showing the positions of the functional groups discussed in the text. **B**. A closeup view showing polar contacts (dashed lines) and hydrophobic interactions (semitransparent spheres) positioning the 18E6 phosphopeptide (cyan sticks) and FSC (thin pink sticks) in the amphipathic groove of a 14-3-3ζ subunit (semitransparent light grey ribbon). $2F_o$-$F_c$ electron density maps contoured at 1σ are shown for the peptide and FSC. # denotes the C-terminus (-COOH). The GGGG linker is omitted for clarity.

FSC occupies its well-defined cavity where it is positioned by hydrophobic interactions with Phe117, Ile166, Ile217 and Leu216, polar contacts with residues Asn42 and Asp213, and a remarkable H-bond involving its 3-metoxy oxygen and the side chain of Lys120 of 14-3-3ζ. The latter contact breaks the Lys120 interaction with the carboxyl-group of the 18E6 PBM formed in the absence of FSC, displacing the carboxyl to another position, where it establishes a new contact with the 12-hydroxy group of FSC. 14-3-3ζ Lys49 also switches its position, and loses a contact to the backbone carbonyl of pThr156 to establish instead a contact with the 12-hydroxy group of FSC. The side chain of the C-terminal Val158 of the 18E6 PBM shifts 3.5 Å towards the phosphate moiety of Thr156, breaking the hydrophobic contact with Val46 of 14-3-3ζ and significantly dispersing the local electron density. As a result, while most of the peptide conformation remained unchanged, the B-factors of the last 18E6 PBM residue in the refined FSC-bound structure increased significantly.

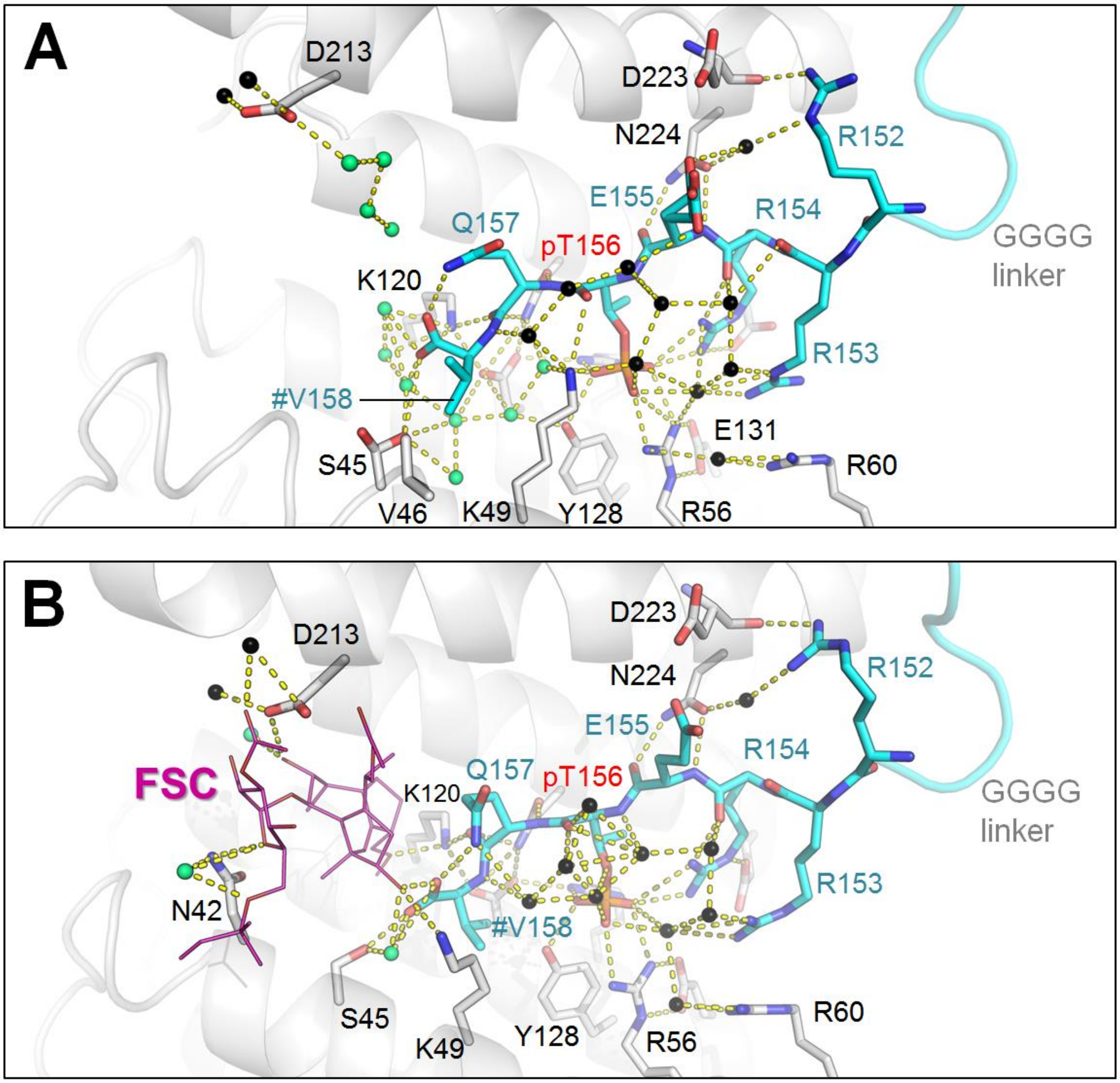


**Supplementary figure 7**. Comparison of the water-mediated polar contacts formed in the 14-3-3ζ/18E6 interface in the absence (**A**) or in the presence of fusicoccin (FSC) (**B**). The main residues involved in the interactions are shown by sticks with color coding: 14-3-3 residues are shown in light grey, 18E6 residues are in cyan, phospho-group of Thr156 is shown by orange sticks. FSC is shown by thin magenta sticks, water molecules affected by FSC binding are shown by lime green, those similar in two structures are black. The C-terminal 18E6 residue (V158) is denoted by #. Note the significant redistribution of water molecules upon FSC binding.

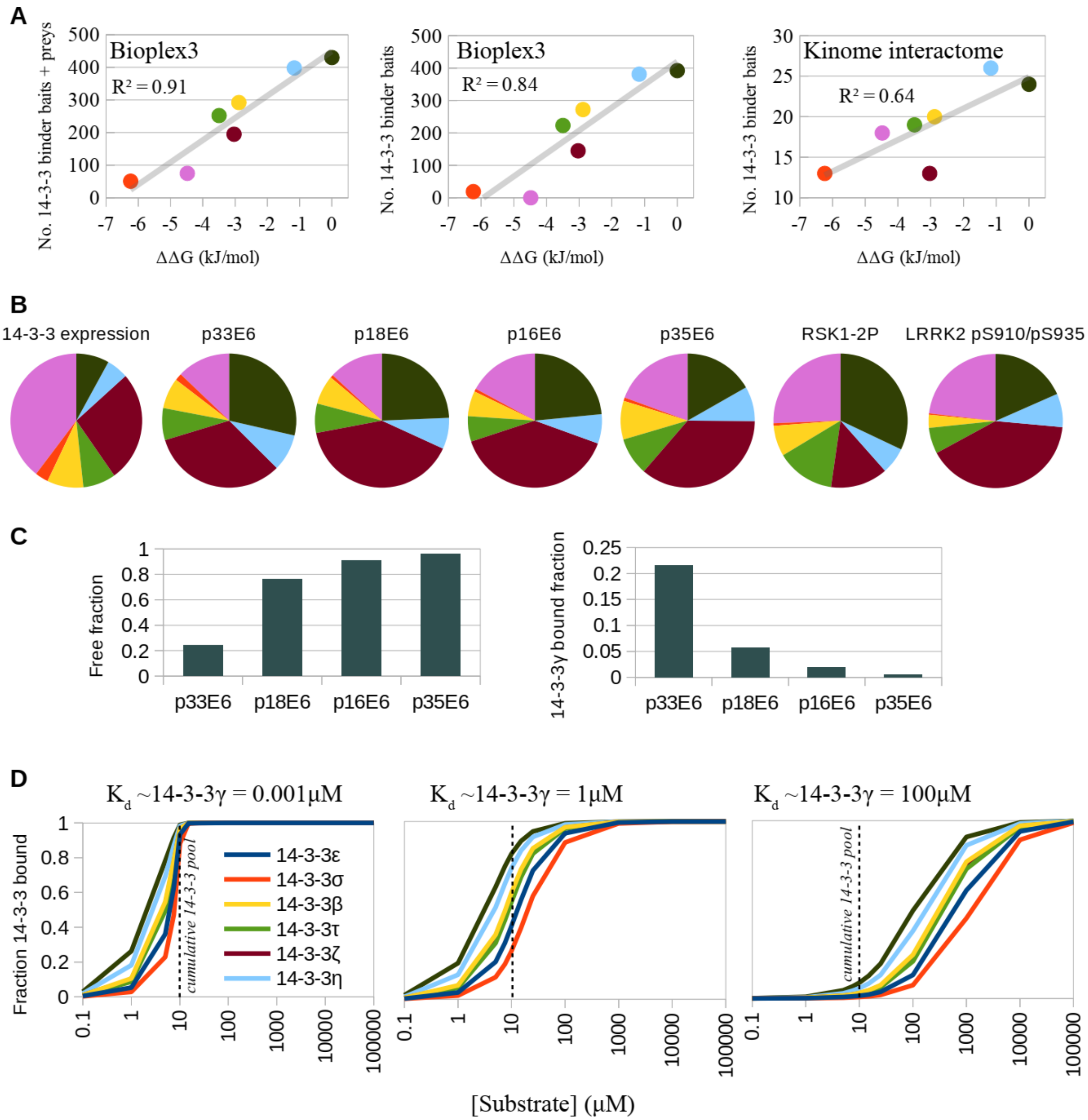


**Supplementary figure 8**. Additional data for Fig. 4. **A**. Correlation between the number of 14-3-3 interaction partners, and the average free binding energy difference ΔΔG between the strongest phosphopeptide-binder 14-3-3γ, and all individual isoforms (same color code as in Fig. 4D). ΔΔG values were calculated from the average $K_D$ ratios from Fig. 5C. Interaction partners can be either "preys" or "baits" in the AP-MS experiment. Using the Bioplex database (https://bioplex.hms.harvard.edu and [18]), the cumulative number of unique interaction partners are shown in the left panel, the number of baits are shown in the middle panel and the number of preys are shown in Fig. 4G. The numbers of 14-3-3 binding baits are shown in the right panel using the kinome interactome as taken from (https://sec-explorer.shinyapps.io/Kinome_interactions/ and [19]). **B**. Predicted proportions of 14-3-3-bound phosphoproteins that would be engaged with each individual isoform are mostly dependent on the proteomic context (see Fig. 4) and less dependent on the absolute affinity of the interaction partner. Predictions were performed using the experimental affinity values, assumed low target concentration (25 nM), and the proteomic context of uterus. Concentrations were calculated from abundancies from the PAXdb (https://pax-db.org and [20]), according to conversion rules described in the Methods section. **C**. The amount of complex formation is strongly dependent on the absolute affinity of the interaction partner. The same prediction is shown as in (B), but the amount of free target, or the formed complex with 14-3-3γ are shown for the four studied E6 proteins. **D**. By varying the concentration and affinity of a 14-3-3 target, it is possible to estimate how much target can sequester and sink the cellular 14-3-3 pool. In the case of a strong binder, an equimolar amount of target is sufficient to saturate the cellular 14-3-3 proteins. However, if the target affinity is relatively weak (>1 μM), the required target concentration can exceed the cumulative 14-3-3 concentration by orders of magnitudes.

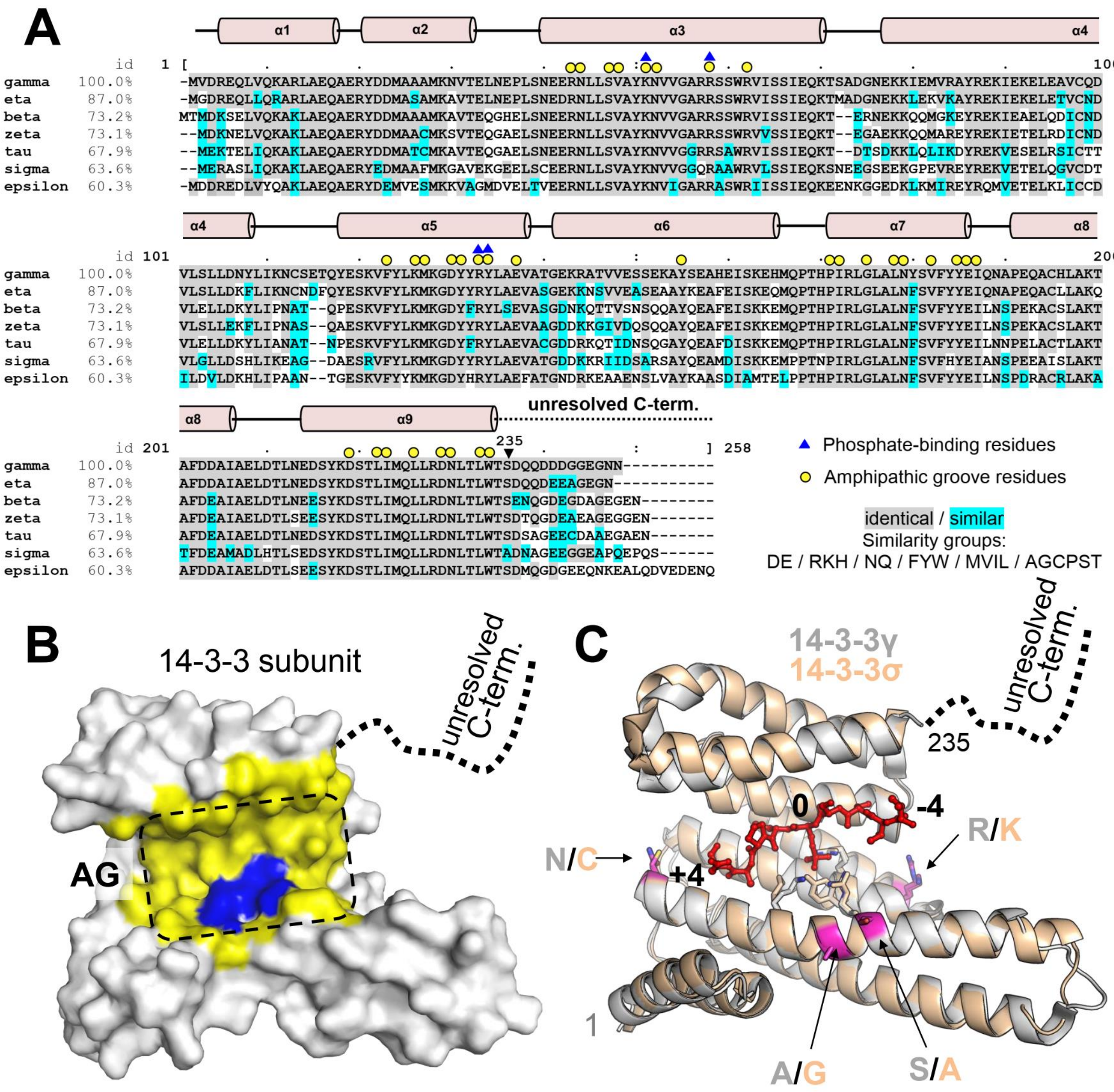


**Supplementary figure 9**. Sequence divergence underlies the phosphopeptide-affinity trend of human 14-3-3 isoforms. **A**. Full-length human 14-3-3 isoforms aligned by Clustal Omega (https://www.ebi.ac.uk/Tools/msa/clustalo/) using 14-3-3γ as a reference show sequence divergence order correlating well with the peptide-affinity trend. Identical residues are highlighted by grey, similar residues are highlighted by cyan (six similarity groups are indicated). Residues forming the phosphopeptide-binding AG groove (yellow surface in panel B) are marked by yellow circles, residues directly coordinating the phosphate group (blue surface in panel B) are marked by blue triangles. Secondary structural elements based on 6A5S structure [21] are shown above the alignment. The variable C-terminal segments are flexible and normally unresolved in crystal structures (dotted line). The last structured residue (S235 in 14-3-3γ) is marked by arrow for clarity. **B**. The phosphopeptide-binding amphipathic groove (yellow) and phosphate-coordinating pocket (blue) are identical in the seven human 14-3-3 isoforms (shown mapped on the surface of 14-3-3σ subunit). **C**. Spatial overlay of the 14-3-3γ (6A5S) [21] and 14-3-3σ (5LU2) [22] structures with the bound phosphopeptide (red sticks), showing the four amino acid differences (magenta) located most close to, yet far enough from the phosphopeptide-binding groove.

**A**

identity level

| | removed: | C | -α3-9,C | -α1α2/ΔC | -α1-α3 | -α1-3/α9 | -α1-3/α9-8 | -α1-3/α9-7 | -α1-3/α9-6 | -α1-3/α9-5 | -α1-5/α9-6 |
|---|---|---|---|---|---|---|---|---|---|---|---|
| | left: | α1-α9 | α1α2 | α3-α9 | α4-α9 | α4-α8 | α4-α7 | α4-α6 | α4α5 | α4 | α6 |
| | **FULL** | **aa1-234** | **aa1-32** | **aa33-234** | **aa71-234** | **aa71-204** | **aa71-185** | **aa71-164** | **aa71-137** | **aa71-116** | **aa139-164** |
| **gamma** | 100,0% | 100,0% | 100,0% | 100,0% | 100,0% | 100,0% | 100,0% | 100,0% | 100,0% | 100,0% | 100,0% |
| **eta** | 87,0% | 87,6% | 84,8% | 88,2% | 86,0% | 82,8% | 81,7% | 78,7% | 83,6% | 76,1% | 65,4% |
| **beta** | 73,2% | 75,7% | 73,5% | 75,9% | 73,5% | 68,2% | 67,3% | 60,9% | 67,7% | 57,1% | 46,2% |
| **zeta** | 73,1% | 76,0% | 75,0% | 75,9% | 74,1% | 69,7% | 69,0% | 63,0% | 69,2% | 59,5% | 42,3% |
| **tau** | 67,9% | 71,7% | 65,6% | 72,4% | 70,4% | 63,6% | 61,9% | 54,3% | 61,5% | 47,6% | 38,5% |
| **sigma** | 63,6% | 66,5% | 62,5% | 67,0% | 65,2% | 60,4% | 60,0% | 54,2% | 59,4% | 47,8% | 35,7% |
| **epsilon** | 60,3% | 64,5% | 60,6% | 64,5% | 62,8% | 56,0% | 53,9% | 46,8% | 50,7% | 38,6% | 34,6% |
| | 1 | 2 | 3 | 4 | 5 | 6 | 7 | 8 | 9 | 10 | 11 |

**B**

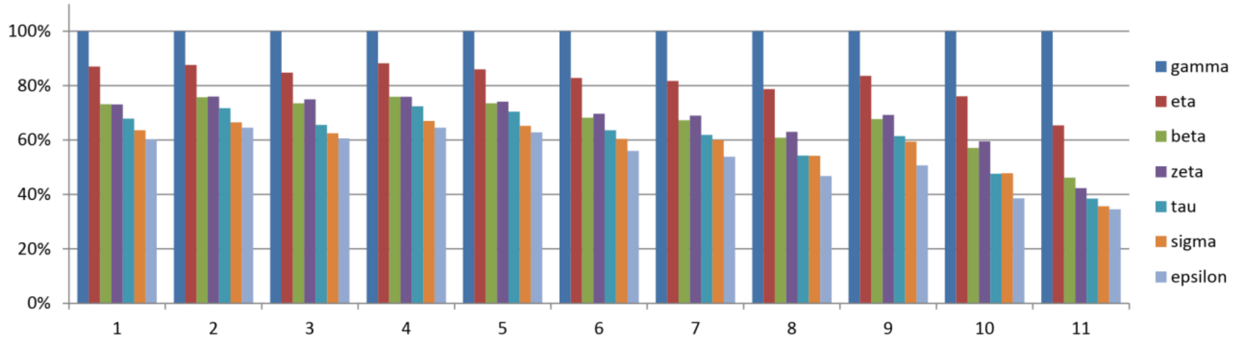


**Supplementary figure 10**. Sequence divergence trend for the seven human 14-3-3 isoforms is preserved throughout the entire 14-3-3 sequence, also for its different sub-regions (indicated from 1 to 11 starting from the full-length sequences). **A.** The identity level relative to 14-3-3γ (the strongest binder) is shown in % and color coded using the standard red-yellow-white scale for different human 14-3-3 isoforms, considering the full-length sequences (1) or different parts thereof (2-11). **B**. The same identity levels as in A shown by histograms for clarity. Note that exclusion of the most variable flexible C-terminal peptides from analysis does not re-shuffle the positions of the 14-3-3 isoforms in the trend significantly. This indicates that the general target affinity differences arise from fine conformational effects spanning the entire structure, rather than a defined sub-region.

#### Supplementary references


1. Wurtele M, Jelich-Ottmann C, Wittinghofer A, Oecking C. Structural view of a fungal toxin acting on a 14-3-3 regulatory complex. *EMBO J* **22**, 987-994 (2003).

2. Ottmann C, *et al.* A structural rationale for selective stabilization of anti-tumor interactions of 14-3-3 proteins by cotylenin A. *J Mol Biol* **386**, 913-919 (2009).

3. Saponaro A, *et al.* Fusicoccin Activates KAT1 Channels by Stabilizing Their Interaction with 14-3-3 Proteins. *Plant Cell* **29**, 2570-2580 (2017).

4. Molzan M, *et al.* Impaired binding of 14-3-3 to C-RAF in Noonan syndrome suggests new approaches in diseases with increased Ras signaling. *Mol Cell Biol* **30**, 4698-4711 (2010).

5. Anders C, *et al.* A semisynthetic fusicoccane stabilizes a protein-protein interaction and enhances the expression of K+ channels at the cell surface. *Chem Biol* **20**, 583-593 (2013).

6. Andrei SA, *et al.* Rationally Designed Semisynthetic Natural Product Analogues for Stabilization of 14-3-3 Protein-Protein Interactions. *Angew Chem Int Ed Engl* **57**, 13470-13474 (2018).

7. De Vries-van Leeuwen IJ, *et al.* Interaction of 14-3-3 proteins with the estrogen receptor alpha F domain provides a drug target interface. *Proc Natl Acad Sci U S A* **110**, 8894-8899 (2013).

8. de Vink PJ, Briels JM, Schrader T, Milroy LG, Brunsveld L, Ottmann C. A Binary Bivalent Supramolecular Assembly Platform Based on Cucurbit[8]uril and Dimeric Adapter Protein 14-3-3. *Angew Chem Int Ed Engl* **56**, 8998-9002 (2017).

9. Sijbesma E, *et al.* Site-Directed Fragment-Based Screening for the Discovery of Protein-Protein Interaction Stabilizers. *J Am Chem Soc* **141**, 3524-3531 (2019).

10. Edwards MR, *et al.* Henipavirus W Proteins Interact with 14-3-3 To Modulate Host Gene Expression. *J Virol* **94**,  (2020).

11. Qin S, *et al.* Structural basis for histone mimicry and hijacking of host proteins by influenza virus protein NS1. *Nat Commun* **5**, 3952 (2014).

12. Gogl G, *et al.* Dual Specificity PDZ- and 14-3-3-Binding Motifs: A Structural and Interactomics Study. *Structure* **28**, 747-759 e743 (2020).

13. Sluchanko NN, Tugaeva KV, Titterington J, Antson AA. Unpublished work (2020).

14. Crooks GE, Hon G, Chandonia JM, Brenner SE. WebLogo: a sequence logo generator. *Genome research* **14**, 1188-1190 (2004).

15. Simon MA, *et al.* High-throughput competitive fluorescence polarization assay reveals functional redundancy in the S100 protein family. *FEBS J* **287**, 2834-2846 (2020).

16. Goldschmidt L, Cooper DR, Derewenda ZS, Eisenberg D. Toward rational protein crystallization: A Web server for the design of crystallizable protein variants. *Protein Sci* **16**, 1569-1576 (2007).

17. Goldschmidt L, Cooper DR, Derewenda ZS, Eisenberg D. SERp Server. (ed^(eds) (2007).

18. Huttlin EL, *et al.* Dual Proteome-scale Networks Reveal Cell-specific Remodeling of the Human Interactome. *bioRxiv*, 2020.2001.2019.905109 (2020).

19. Buljan M, *et al.* Kinase Interaction Network Expands Functional and Disease Roles of Human Kinases. *Molecular Cell* **79**, 504-520.e509 (2020).

20. Wang M, Herrmann CJ, Simonovic M, Szklarczyk D, von Mering C. Version 4.0 of PaxDb: Protein abundance data, integrated across model organisms, tissues, and cell-lines. *Proteomics* **15**, 3163-3168 (2015).

21. Xu Y, Ren J, He X, Chen H, Wei T, Feng W. YWHA/14-3-3 proteins recognize phosphorylated TFEB by a noncanonical mode for controlling TFEB cytoplasmic localization. *Autophagy* **15**, 1017-1030 (2019).

22. Sluchanko NN, *et al.* Structural Basis for the Interaction of a Human Small Heat Shock Protein with the 14-3-3 Universal Signaling Regulator. *Structure* **25**, 305-316 (2017).
